# Extracellular vesicles are the main contributor to the non-viral protected extracellular sequence space

**DOI:** 10.1101/2023.08.07.552246

**Authors:** Dominik Lücking, Coraline Mercier, Tomas Alarcón-Schumacher, Susanne Erdmann

**Affiliations:** Max-Planck Institute for Marine Microbiology, Celsiusstraße 1, 28359 Bremen, Germany

**Keywords:** Viromics, metagenomics, eDNA, Extracellular vesicles, Genome transfer agents

## Abstract

Marine environmental virus metagenomes, commonly referred to as ’viromes’, are typically generated by physically separating virus-like particles (VLPs) from the microbial fraction based on their size and mass. However, most methods used to purify VLPs, enrich extracellular vesicles (EVs) and gene transfer agents (GTAs) simultaneously. Consequently, the sequence space traditionally referred to as a ’virome’ contains host-associated sequences, transported via EVs or GTAs. We therefore propose to call the genetic material isolated from size-fractionated (0.22 µm) and DNase-treated samples *protected environmental DNA (peDNA).* This sequence space contains viral genomes, DNA transduced by viruses and DNA transported in EVs and GTAs. Since there is no genetic signature for peDNA transported in EVs, GTAs and virus particles, we rely on the successful removal of contaminating remaining cellular and free DNA when analyzing peDNA. Using marine samples collected from the North Sea, we generated a thoroughly purified peDNA dataset and developed a bioinformatic pipeline to determine the potential origin of the purified DNA. This pipeline was applied to our dataset as well as existing global marine ’viromes’. Through this pipeline, we identified known GTA and EV producers, as well as organisms with actively transducing proviruses as the source of the peDNA, thus confirming the reliability of our approach. Additionally, we identified novel and widespread EV producers, and found quantitative evidence suggesting that EV-mediated gene transfer plays a significant role in driving horizontal gene transfer (HGT) in the world’s oceans.

## Introduction

The presence of extracellular entities strongly shapes microbial communities. Particles of various origins mediate the transport of genetic material from one cell to another, thus playing a crucial role in horizontal gene transfer (HGT) (1, 2). Most prominently, viruses are highly abundant and diverse drivers of ecological and evolutionary interactions within a community (3, 4, 5, 6). However, due to the limited culturability of their hosts, viruses often escape traditional culture-based approaches (7), leading to the development of culture-independent techniques to study their fundamental impact on microbial communities, global biogeochemical cycles and their effect on climate change. Similar to metagenomic studies, researchers have sequenced and analyzed the genetic content of the viral fraction on a community level, leading to the advent of viral metagenomics or ‘viromics’ (8). This approach traditionally relies on the physical, pre-sequencing separation of viruses-like particles (VLPs) from microbial cells. Methods like sequential size filtration, ultracentrifugation, tangential flow filtration, and flow cytometry exploit the distinct physical properties of VLPs when compared to microbes (9, 10, 11, 12). Additionally, bioinformatic methods were developed to identify virus-like sequences among microbial sequences (13, 14, 15, 16). This led to the discovery of many diverse viruses, fulfilling crucial functions in their respective microbial community (17). Interestingly, even after the most thorough removal of microbial cells, non-viral genes were shown to be present in ‘viromes’ generated from many diverse environments (7, 18). The contamination of ‘viromes’ with microbial sequences originating from remaining intact cells and free extracellular DNA has been reported in several studies. Consequently, tools have been developed to estimate the proportion of true viral DNA in a given virome, using the abundance of reads with homologs in available prokaryotic databases or a specific set of microbial marker genes (e.g. 16S rRNA gene) (18, 19, 20). These tools fall short of assessing the true degree of contamination of a ’virome’ , because the abundance of prokaryotic-, non-virus-like genes is not necessary due to microbial contaminations, but can be the result of horizontal gene transfer processes.

Long before the development of modern ‘viromics’, studies showed that viruses carry and distribute random microbial genes (21, 22) or specific ‘auxiliary metabolic genes’ (AMGs) in addition to bona fide viral genes (e.g. genes necessary for particle assembly or viral genome replication), in a well-described process termed ‘transduction’. Here, either genetic material adjacent to the integrated viral genome (specialized transduction) or random snippets of the host genome (general transduction) are packaged into the viral particles (1). AMGs have been shown to fundamentally alter the metabolism of microbes by providing genes otherwise unavailable to their host (8, 23), further demonstrating the need for a good understanding of non-viral DNA in viromes.

Viromes are traditionally generated by separating VLPs from cells. However, methods that enrich VLPs by removing larger and heavier microbial cells also enrich entities similar in size and mass to VLPs. Most prominently, gene transfer agents (GTAs) and extracellular vesicles (EVs, also referred to as membrane vesicles, MVs, or outer membrane vesicles, OMVs) are particles with similar physical properties and both have been shown to be involved in HGT, thus contributing to the presence of non-viral DNA in ‘viromes’.

GTAs are particles transporting host DNA from one cell to another. They likely derived from defective prophages, and retained functional genes for the head and tail components of a head-tailed virus particle, including the genes for DNA packaging. Therefore, mass and size (40 - 60 µm) of GTA particles are very similar to head-tailed viruses, making it hard to differentiate them from viruses solely based on morphology. Notably and in contrast to true viruses, GTAs do not specifically package the GTA-producing gene cluster into the particle, but transport short segments of the host genome. Up to this date, several distinct gene clusters have been identified that produce GTAs (24, 25, 26, 27, 28).

Prokaryotic EVs are small (10 - 300 nm) spherical structures derived from the cell membrane (29). EVs represent compartments that protect their cargo from degradation and are used for the transport of a variety of different components across the extracellular space. This includes the transport of nutrients, toxins, antigens, lipids, proteins, RNA, and DNA (30, 31, 32, 33, 34, 35, 36). Recent studies showed an abundance of EVs in marine environments of up to 10^6 vesicles per milliliter (33), produced across diverse taxa. EVs, produced by highly abundant marine heterotrophs and autotrophs, such as Pelagibacter, Marinobacter, and Prochlorococcus, have been shown to transport fragments of chromosomal and plasmid DNA (37, 38), thus contributing to the fraction of non-viral DNA within viromes.

In this study, we aimed to explore the non-viral sequence space of viromics datasets. First, we generated our own dataset, carefully avoiding possible contaminations. Then we categorized the sequences from this dataset and publicly available viromics datasets as virus- or non-virus-derived. Subsequently, we explore the non-virus-derived sequence space to detect the extent of non-viral DNA potentially being horizontally transferred between cells. We then identify the means of transport (GTA-, EV-, or virus-driven) for the sequences by linking the datasets to existing microbial metagenomes and genomes. We identify potential novel EV- and GTA producers and metagenomics-assembled genomes (MAGs) with an actively transducing virus. We propose using the term ‘protected extracellular DNA’ (peDNA) for DNA sequence data derived from appropriately purified environmental fractions <0.02 µm, so far referred to as viromics datasets.

## Results and Discussion

### Viromics datasets represent the sequence space of protected extracellular DNA (peDNA)

The majority of samples prepared for ‘viromics’ include GTAs and EVs, in addition to virus particles. All three entities are small protein- and/or lipid-containing particles that can enclose cellular DNA (31) or were found to bind cellular DNA on their surface (39), thus inflating the sequence space that traditionally has been described as a ‘virome’. In contrast to free extracellular DNA, DNA that is enclosed in or tightly associated with particles is protected against degradation by extracellular nucleases occurring in the environment or nucleases used to clean samples from free extracellular DNA. Hence, we propose the term ‘protected extracellular DNA’ (peDNA) to describe the entirety of DNA transported by viruses, GTAs and EVs (Figure 1). We will use this term throughout this work.

**Figure 1:**
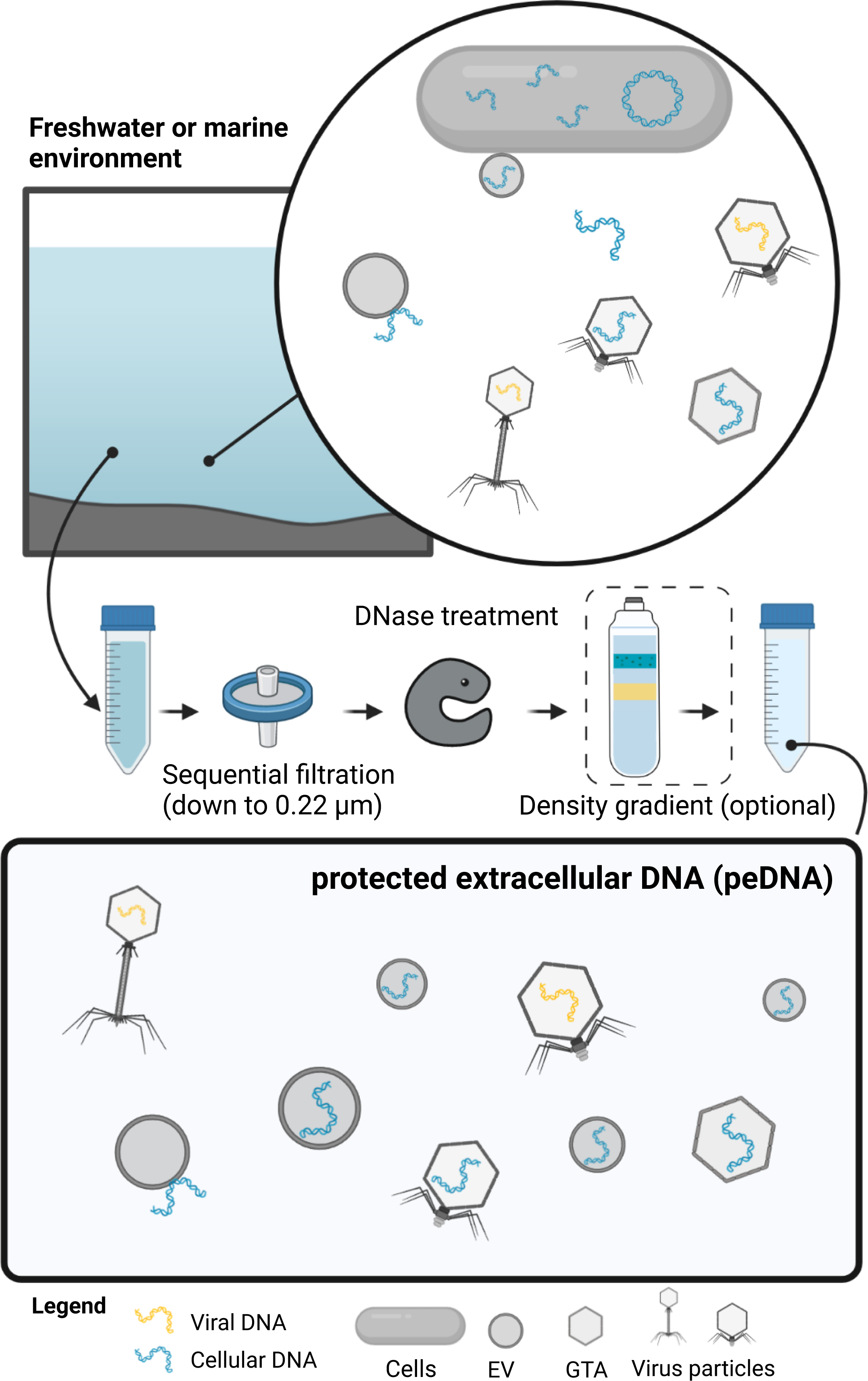
Conceptual composition of protected extracellular DNA. The top panel depicts microbial entities present in a water body: microbial cells, viruses containing viral and microbial genetic material, gene transfer agents and extracellular vesicles containing host DNA. After size filtration (0.22 µm) and DNase treatment and, if applicable, purification via density gradients, microbial cells and free DNA are removed (middle panel). The remaining DNA makes up the sequence space of protected extracellular DNA, peDNA (bottom panel).

### Purification of environmental samples for the generation of peDNA datasets is essential to explore of the entire dataset

Previously, the percentage of 16S/18S rRNA-mapping reads was used as a proxy for host contamination in virome datasets (19). However, it has since been shown that GTAs and EVs enclose host DNA randomly, including 16S/18S rRNA genes (40). We calculated SSU rRNA alignment rates for two highly purified samples: DNA extracted from virus isolates, purified by sequential plaque assays and 0.2 um size filtration (41) and DNA extracted from EVs purified from culture supernatants of *Prochlorococcus* (33) (Figure 2). While the alignment rates were low for virus isolates (mean = 0.000437 %), the percentage of 16S/18S-mapping reads was five orders of magnitude higher in DNA extracted from purified EVs (1.81 %), even exceeding the mean alignment rate of publicly available microbial environmental metagenomes (mean = 0.078 %, (19)). For these samples, Biller *et al* confirmed the absence of microbial cells using electron microscopy. Thus, the presence of 16S/18S rRNA-mapping reads neither proves or disproves contamination in peDNA samples. The only way to exclude contaminations with cellular DNA or extracellular free DNA is the rigorous purification of the sample before sequencing.

**Figure 2:**
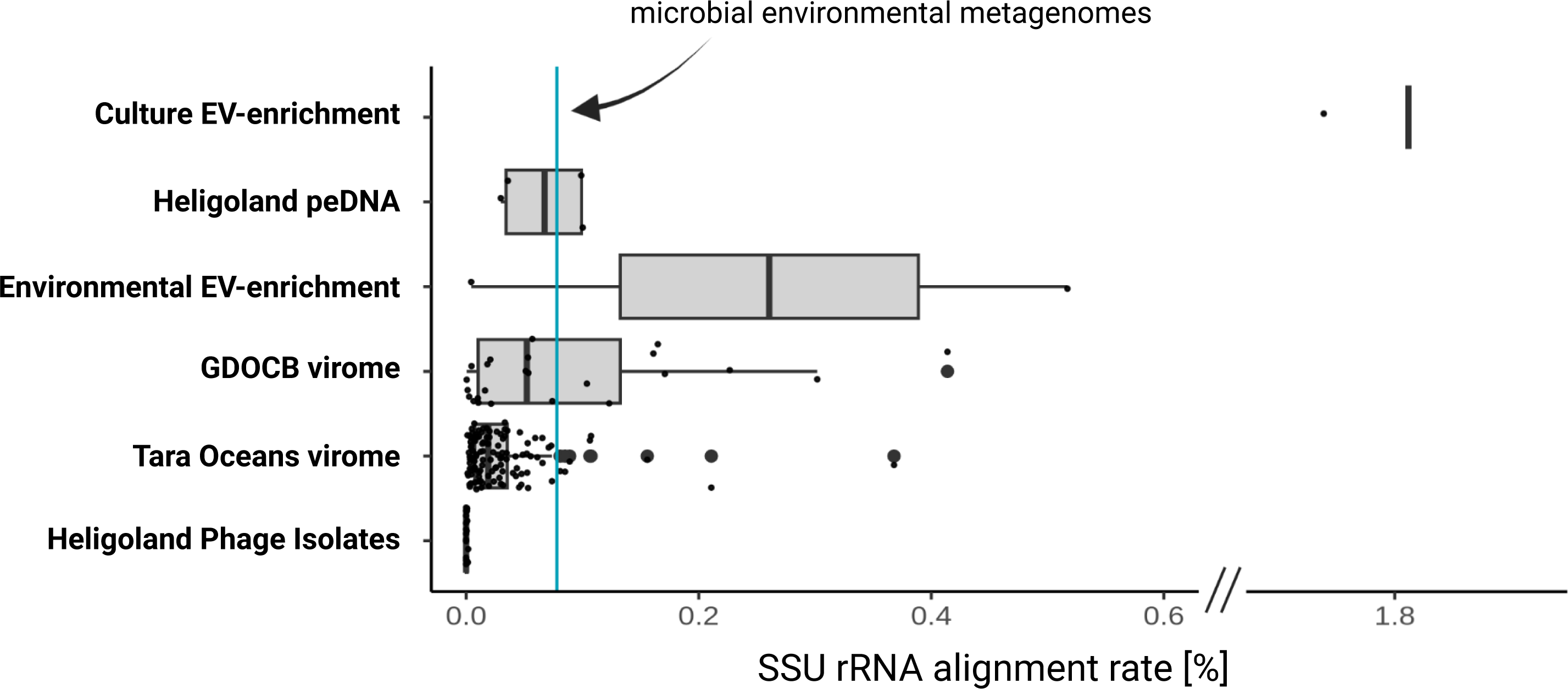
Comparison of SSU rRNA alignment rates of diverse viromes and EV preparations. Each dot represents the percentage of reads aligning to either 16S or 18S rRNA genes. The cyan line indicates the average alignment rate for publicly available metagenomes from various environments (19). ‘Culture EV enrichment’: a cell free preparation of EVs from *Prochlorococcus* cultures (33). ‘Heligoland peDNA’: dataset generated in this study from a highly purified (filtration, DNase treatment, gradient purification) <0.2 µm Heligoland water fraction. ‘Environmental EV enrichment’: Density gradient-purified EVs isolated from seawater samples (33). ‘GDOCB virome’: <0.2 µm fraction enriched for VLPs by flow cytometry (42). ‘Tara Oceans virome’: <0.2 µm fraction purified by size filtrations and DNase treatment (17). ‘Heligoland phage Isolates’: DNA extracted from virus isolates, purified by 0.2 µm size filtration (41).

For this purpose, we generated a dataset from rigorously purified samples using several sequential filtration steps, DNase treatment and density gradient purification (Methods, Supplementary Figure 1), resulting in cell-free samples containing virus-like particles, GTA particles and EVs. Subsequently, we compared the SSU rRNA alignment rates of our dataset with one metagenomic peDNA dataset and two viromic datasets: Density gradient-purified EVs isolated from seawater samples (’Environmental EV enrichment’) (33); the ’Tara Oceans virome’ dataset (17), purified by size filtration and DNase treatment; and the ’GDOCB virome’ dataset (42). GDOCB viromes are purified by flow cytometry. The process excludes free DNA and microbial cells with physical properties outside of the analyzed size spectrum, which makes it possible to assume that these viromes are free from contamination. Both datasets showed increased SSU rRNA alignment rates (Figure 2), with some samples even exceeding the microbial metagenome alignment rate of 0.078 %. Likewise, our dataset, while on average showing lower alignment rates (mean = 0.066 %), contained samples exceeding that threshold. Lastly, Tara Oceans viromes mostly showed very low SSU rRNA alignment rates, with very few exceptions (mean = 0.031 %). Overall, even thoroughly purified and confirmed contamination-free datasets show highly variable SSU rRNA alignment rates. We concluded that the majority of SSU rRNA hits in these datasets are enclosed in VLPs, GTA’s or EVs rather than in contaminating microbial cells, and therefore included the datasets in the subsequent analysis.

### Separation of non-viral (nvpeDNA) from viral protected extracellular DNA (vpeDNA) indicates that EVs and GTAs could be very abundant entities in the ocean

The sequence space of protected extracellular DNA (peDNA) consists of virus genomes (viral protected extracellular DNA, vpeDNA) and non-viral, microbial DNA (non-viral extracellular DNA, nvpeDNA), deriving from transducing viruses, GTAs and EVs. nvpeDNA represents the sequence space that is potentially horizontally transferred between cells and therefore has major implications on the ecology and evolution of the organism in this environment and the environment itself.

In order to separate non-viral peDNA from viral peDNA, we developed a bioinformatic pipeline that, in brief, identifies virus sequences, separates those from non-viral sequences and calculates a non-viral to viral peDNA ratio in a given dataset (Figure S1 and Methods). This pipeline was first applied to isolated viruses and purified EVs (Figure 3A). As expected, vpeDNA made up >99 % of the DNA of purified Heligoland phage isolates and nvpeDNA made up 98 % of the DNA transported in purified EVs of a pure *Prochlorococcus* culture, verifying that the pipeline is reliably separating vpeDNA from nvpeDNA.

**Figure 3:**
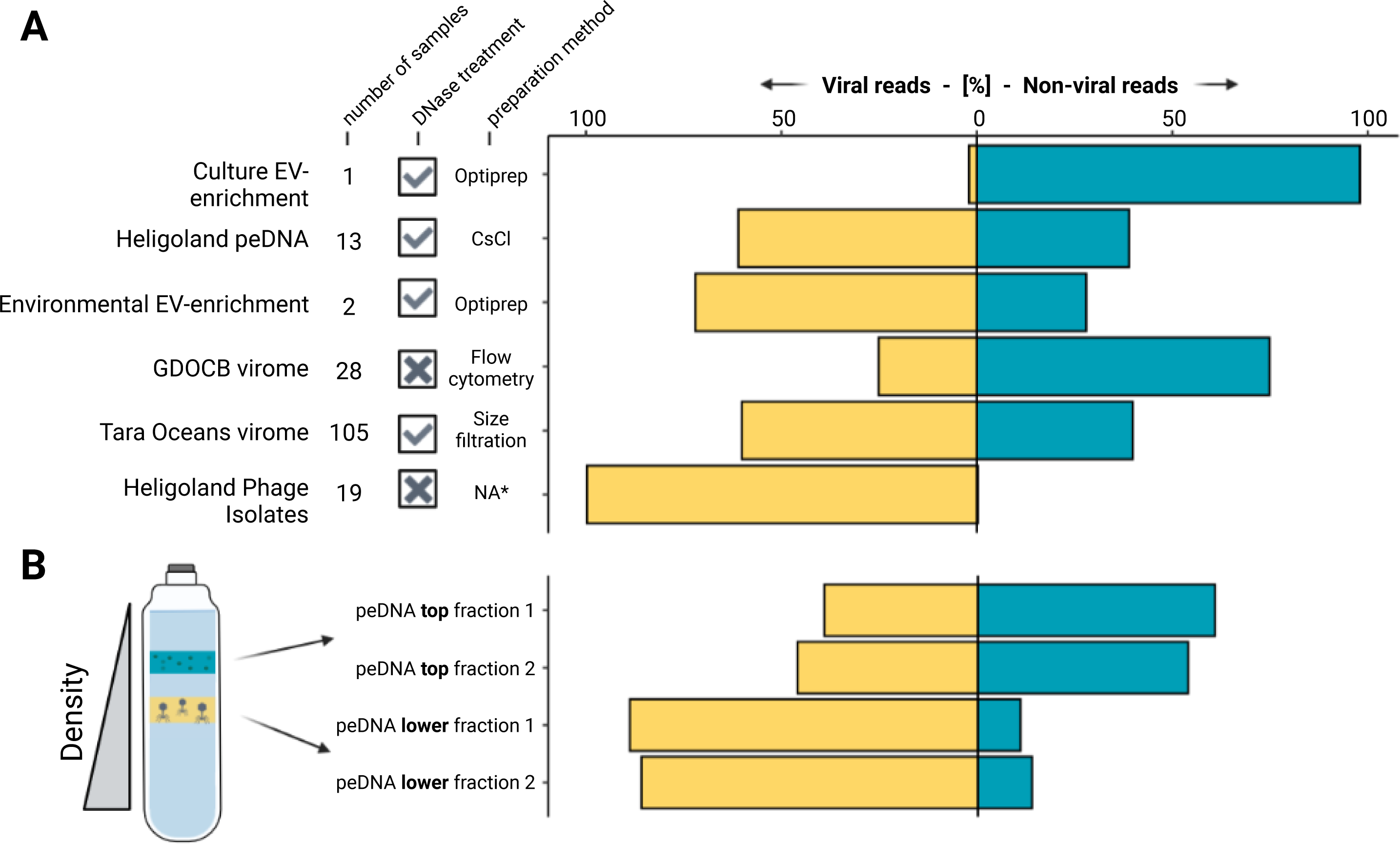
**(A) Non-viral to viral peDNA ratio across different studies.** Each bar represents the percentage of read pairs mapping to contigs classified to be non-viral or viral for viromes, peDNA enrichments, EV enrichments and pure phage isolates. Sample size and purification methods are indicated for each sample. (**B) Non-viral to viral peDNA ratio in different fractions of CsCl gradients.** Left, schematic view of seawater samples running through CsCl gradients (adapted from 43). Right, non-viral to viral peDNA ratio for top and bottom fractions in CsCl gradients for the Heligoland peDNA sample.

Consequently, we applied the pipeline to the entire Helgoland peDNA dataset and the other datasets that we verified earlier to be reasonably contamination-free peDNA datasets (Figure 3A). Helgoland peDNA contained 39 % non-viral reads. In the Tara Oceans Viromes, nvpeDNA made up, on average, 40 % of all reads (105 samples). While viruses are considered the most abundant nucleic acid-containing biological entities in the ocean (3), these findings clearly indicate that EVs and GTAs, transferring cellular DNA, are likely very abundant entities as well. Surprisingly, in GDOCB viromes, the proportion of nvpeDNA to vpeDNA was even higher (75 % nvpeDNA). However, this may be due to the comparably low sequencing depth, resulting in incomplete assembly and therefore hindering a reliable identification of virus contigs. Additionally, since the samples were not DNase treated, particles with the same size as the sorted VLPs might carry free DNA attached to the surface of the particles, therefore inflating nvpeDNA. In order to avoid any artificially introduced biases, this dataset was excluded from further downstream work.

We analyzed individual fractions of density gradients that were used to purify the Heligoland peDNA, because it was shown previously that density gradients can separate VLPs from EVs. VLPs, also including GTAs, were found to be more abundant in lower fractions of the gradients, while EV-like particles were more abundant in upper fractions (33, 43). Indeed, in the upper gradient fractions of the Helgoland peDNA, nvpeDNA made up 60 % and 54 % of all reads in two biological replicates, while in the lower fraction nvpeDNA made up only 14 % and 11 % (Figure 3B), confirming the previous observations. Thus the proportion of nvpeDNA is much higher in the upper fraction, that enriches EVs additionally to some VLPs, compared to the lower fraction, that enriches mainly virus particles and GTA’s. This indicates that EVs are likely contributing significantly more to the nvpeDNA sequence space than GTA’s and viruses.

### Identifying the origin of non-viral protected extracellular DNA reveals that EVs could be the main driver of horizontal gene transfer in the oceans

For contamination-free peDNA samples, we consider three possible origins of nvpeDNA: DNA transduced by viruses, DNA transported in GTA particles and DNA associated with EVs (Figure 4A).

**Figure 4:**
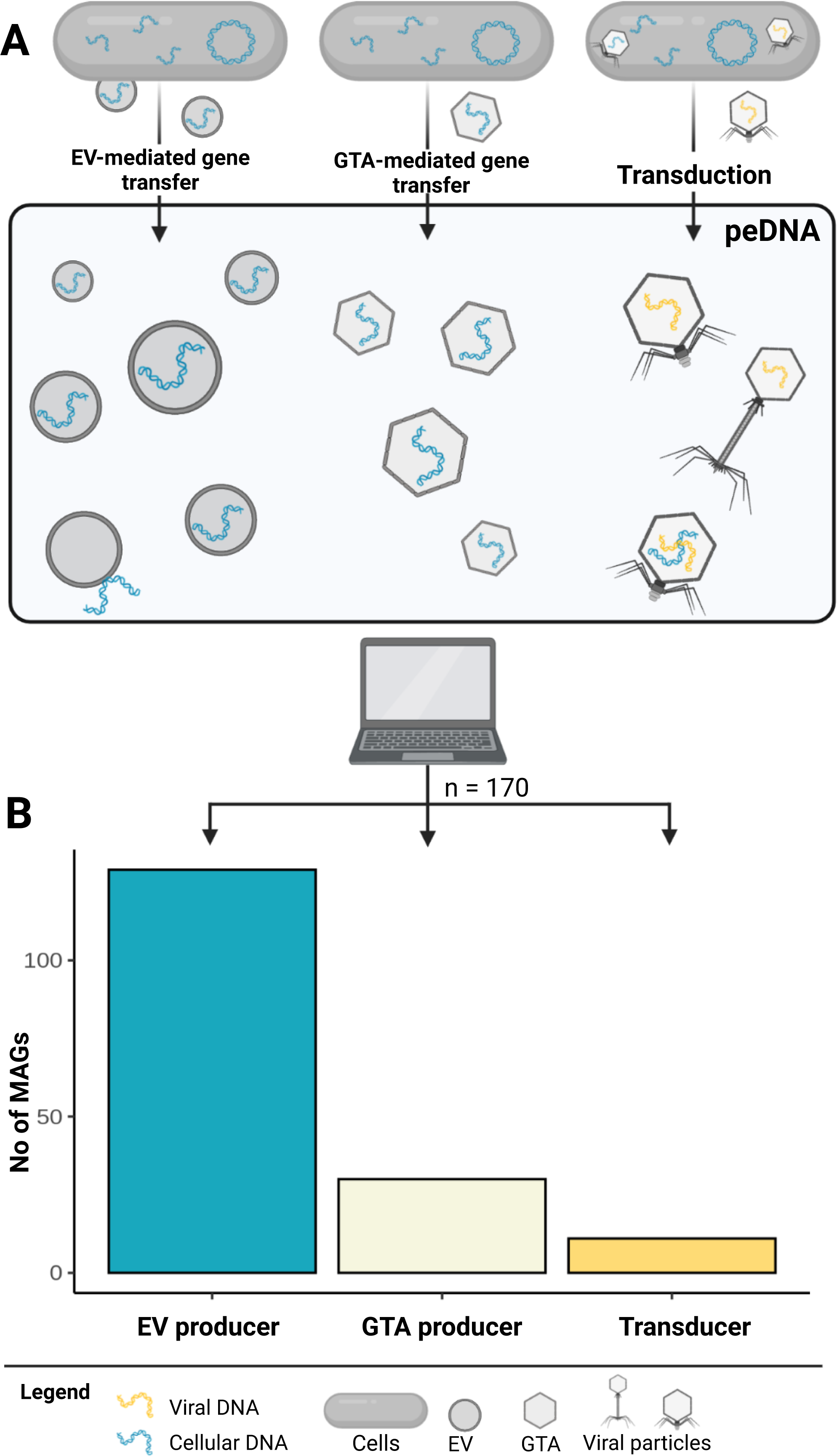
**(A) Mechanisms of horizontal gene transfer that contribute to peDNA.** The origin of sequences comprising peDNA can be either EV-mediated or GTA-mediated gene transfer or via transduction. (**B) Number of MAGs assigned to each mechanism.** Of 200 analyzed metagenomic assembled MAGs, 170 could be assigned to predominantly use one of the three mechanisms in order to transport their genetic material into the extracellular space: 129 EV producers, 30 GTA producers and 11 genomes, with an actively transducing phage.

It is inherently difficult to differentiate the origin of nvpeDNA based on sequence content. To this date, there are no reports on specific sequence signatures (e.g. marker genes) for DNA transported in EVs or GTAs, making DNA transported in EVs indistinguishable from DNA transported in GTA’s or virus particles. Therefore, we developed a bioinformatic approach that tackles this differentiation from a different perspective. First, each read in the nvpeDNA fraction was linked to a given potential microbial host (Figure S2). Then, the 20 most nvpeDNA recruiting MAGs per sample were selected. The main mechanism, which is most likely used to transport its DNA into the extracellular space was predicted, thus linking each read to either EV-, GTA- or transduction-associated transport. We confirmed that the abundance of these organisms (MAGs) in peDNA datasets does not correlate (R² < 0.01) with their abundance in the corresponding metagenomes (Figure S3), and that none of the organisms identified are known to produce particularly small cells that could pass 0.2 µm filters. This additionally indicates that the high abundance of their genomic DNA in the nvpeDNA fraction is not due to cellular contamination with cells, but indeed the active transport of genomic DNA into the extracellular space via EVs, GTAs or virus particles. This approach was applied to nine Tara Oceans viromes which were linked to 2,307 MAGs (44) and the entire Heligoland peDNA dataset which was linked to 457 MAGs sequenced from seawater coming from the same sampling station on Heligoland (45). This yielded 200 MAGs (180 from Tara Oceans plus 20 from Heligoland), subsequently categorized as either GTA- or EV-producer or containing an actively transducing provirus. For details on the categorization approach, refer to the following sections, in brief: MAGs that contained an active (increased coverage in provirus region) provirus were labeled as ‘transducer’. Similarly, the respective MAG was labeled as’ GTA producer’ if a complete or nearly complete GTA cluster could be identified. If neither an active provirus or a GTA cluster was identified, the MAG was labeled as ‘EV producer’ (Figure S2, Figure S4). All labels were manually checked and verified by scrutinizing coverage plots (Figure S5), prophage regions and GTA clusters.

Among the 200 top peDNA-recruiting MAGs 170 could be assigned unambiguously to one of the three categories (Figure 4B). Most importantly, the majority was identified as EV producers, confirming that EVs that have been shown to be very abundant in the marine environment (38), are not just abundant entities but also significantly contribute to the peDNA sequence space and thereby are likely one of the most important drivers of horizontal gene transfer.

### Identification of four novel GTA producers with RcGTA-like clusters

Of the 170 unambiguously assigned MAGs, 30 were identified to contain a functional (>10 GTA-associated genes, core genes present) GTA cluster. For most identified GTA producers, the presence of a GTA cluster has been described elsewhere: *Roseobacter sp*. (n = 16), *Sulfitobacter sp*. (n = 8) and *Roseovarius sp*. (n = 1) are known GTA producers of the order *Rhodobacterales* (46, 47, 48), indicating the efficiency of our approach. Additionally, we detected a functional GTA cluster in four more species: *Tateyamaria sp.*, *Pseudooceanicola sp.*, *Maritimibacter sp.* and *Flavimaricola sp*. all contained RcGTA homologs, including genes encoding the major capsid protein, a terminase, a proteinase and proteins associated with tail assembly (Figure 5). The GTA cluster on the genome of the *Tateyamaria* species was distributed more widely over the genome in partial subclusters, as it has been described elsewhere (26). As far as we know, these four species’ clusters are not described elsewhere. We suggest that they are complete and functional because the core genes necessary to form GTA particles are present (49); however, laboratory experiments are necessary to confirm their full functionality.

**Figure 5:**
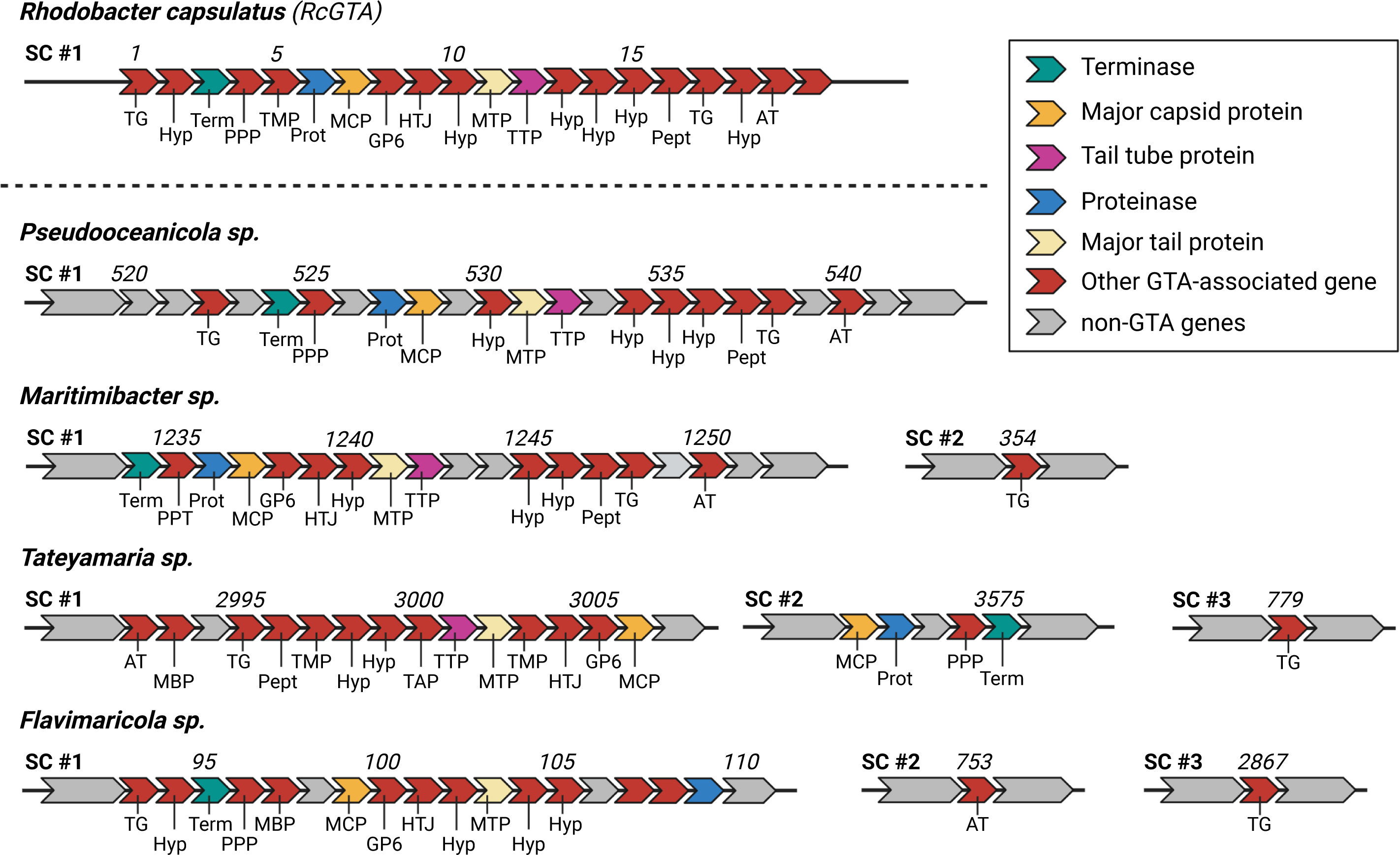
Genome maps of four novel GTA producers. Organization of genes for the four identified, potentially novel GTAs, compared to the GTA cluster of *Rhodobacter capsulatus*. ORF number is given above the map, encoded protein function is indicated below. Non-GTA encoding genes are gray, GTA-associated genes are shown in red. In addition, core GTA encoding genes are colored accordingly. Encoding protein function is given below: *AT -* acetyltransferase, *MBP -* membrane bound protein, *TG* - transglycosylase, *Pept -* phage cell wall peptidase, *TMP -* transmembrane protein, *Hyp -* hypothetical, *TAP -* tail assembly protein, *TTP -* phage tail tube protein, *MTP* - phage major tail protein, *HTJ -* head-tail joining protein, *GP6 -* gp6-like protein, *MCP -* major capsid protein, *Prot -* proteinase, *PPP -* phage portal protein.

### Only a few transducing proviruses could be identified with confidence in the peDNA sequence space

In order to label a MAG as containing an actively transducing provirus and therefore as a transducer, we relied on a combination of virus prediction tools, manual analysis of coverage plots and functional annotation of the proviral regions (see Methods - Identification of potential transducers extracellular vesicle- and gene transfer agent producers). The virus genome itself is present in all viral particles produced, in contrast to the transduced microbial DNA, which could be a randomly selected host DNA fragment (general transduction) or a specifically selected region (specialized transduction). In both cases, the coverage over the proviral region should be increased compared to the surrounding non-viral regions (Figure 6C). Thus, only MAGs which showed the expected coverage profiles were labeled as transducers. Contrastingly, if a region recruited no reads from the peDNA fraction, the region was considered absent and the MAG was labeled as an EV producer (compare Figure 6A *Haliea sp.* Station 158 SRF). In some (17 out of 200) cases, a clear assignment was not possible due to inconclusive coverage profiles or contradicting GTA and prophage predictions. These MAGs were labeled ’unclear’ and removed from further analysis. We identified 11 MAGs that carried an integrated and actively transducing provirus. Interestingly, a *Haliea sp*. with a proviral region was identified that recruited peDNA reads coming from one sampling station but none from the other. However, the non-viral part of the genome recruited high amounts of reads in both stations, albeit with lower coverage in the station where the provirus was absent (Figure 6B). We hypothesize that there are two separate, distinct populations of the same *Haliea* species at the two stations: one, with the provirus integrated and actively transducing and one without the provirus. The fact that DNA from the population without the provirus is present in the peDNA fraction indicates that *Haliea sp.* transports its DNA into the extracellular space differently. In the absence of a GTA cluster, we hypothesize that *Haliea* sp. is capable of EV production and EV-mediated gene transfer. This demonstrates that transduction and EV-mediated gene transfer are not exclusive mechanisms of HGT but can overlap.

**Figure 6:**
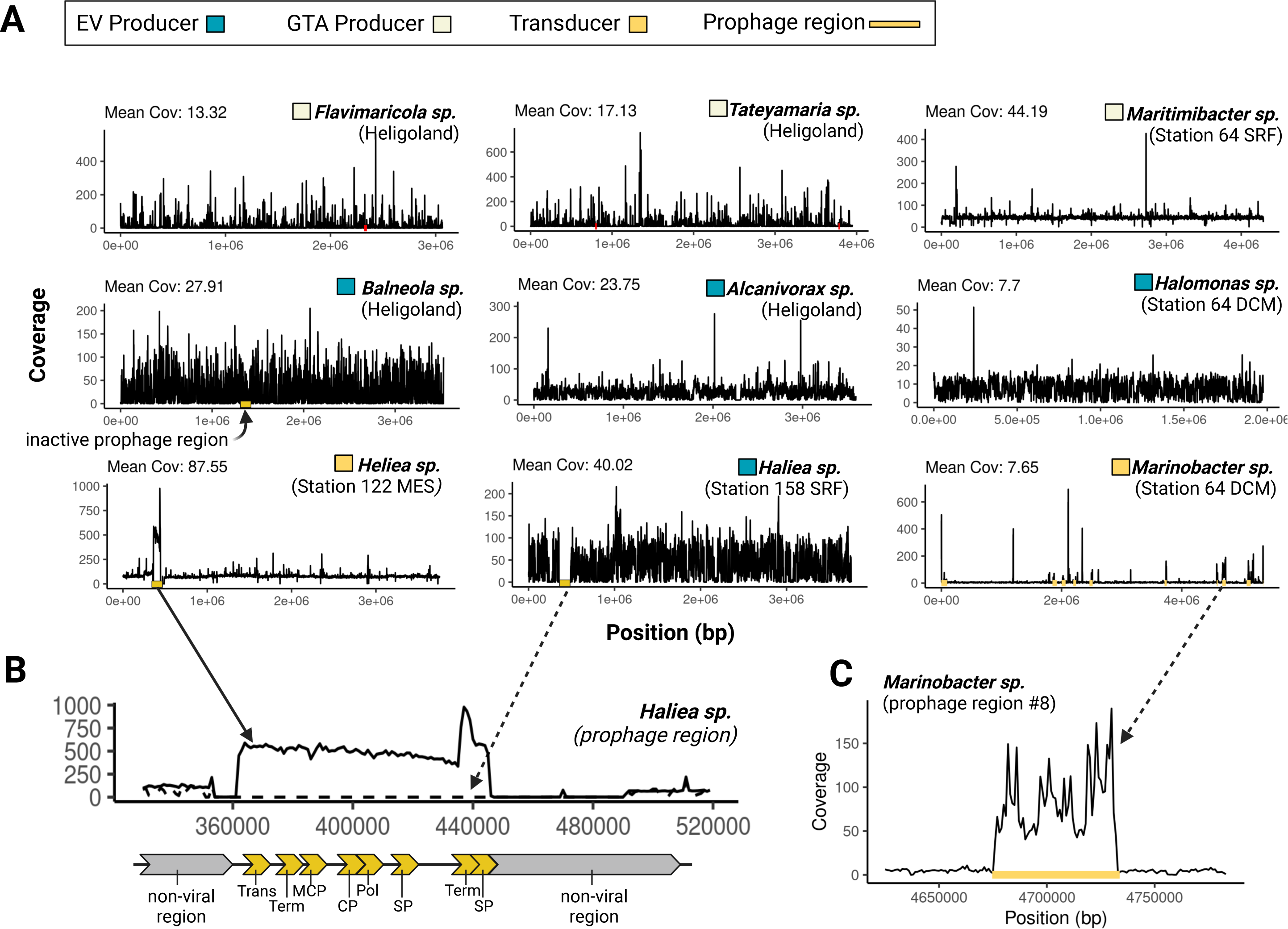
**(A) Coverage plots of identified GTA- and EV-Producers and MAGs containing an actively transducing phage (transducer).** Coverage plots of 4 EV-producer, 2 genomes with an actively transducing phage and 3 GTA-producer (coverage blots of all analyzed MAGs see Figure S12). Active prophage regions are indicated with yellow bars. (**B) Genome map and detailed coverage plot of identified prophage region.** On top, detailed coverage plot of the prophage region in two different samples. Coverage of reads from 122_MES (solid line) and Station 158_SRF (dotted) differs for this specific region. Below, schematic genome map for the genes identified in the prophage region and their approximate positions. *Trans* - transposase, *Term* - terminase, *MCP -* major capsid protein, *CP -* coat protein, *Pol -* Polymerase, *SP -* shaft protein. (**C) Coverage plot of an actively transducing phage.** Close up of the coverage of prophage region #8 of *Marinobacter sp.* and surrounding non-viral regions.

### Identification of known and novel EV producers reveals that EV production is common amongst abundant marine bacteria

We identified 129 MAGs as EV producers. Most identified genera are known EV producers: *Marinobacter* (n = 19), *Alcanivorax* (18), *Flavobacteria* (9), *Thalassospira* (8), *Rheinheimera.* (6) and *Polaribacter* (3), are known to produce high amounts of EVs (38, 50). The fact that most organisms we labeled as EV producers are already known EV producers again supports the efficiency of the approach. Interestingly, multiple MAGs of the genera *Haliea* (n = 16) and *Idiomarina* (n = 8) were identified to be EV producers. So far, EV production by either genus has not been described elsewhere and experimental confirmation is needed. However, this supports the observation by Biller et al, that many marine heterotrophs are actively producing EVs. While EVs could be used as a nutrient source (33, 38), or facilitate horizontal gene transfer (HGT) within microbial communities (37) potentially contributing to the evolution and adaptation of marine microbial populations, future research should aim to clarify the ecological and evolutionary role of EVs in the ocean.

### The functional profile of peDNA links EV production to transposon induced gene mobilization

The functional profile of ‘viromes’ or peDNA, has been assessed previously (51, 20). However, since these studies analyzed this sequence space from a virus perspective, mainly focusing on auxiliary metabolic genes (AMGs), they often excluded genes not directly associated with viral genomes. Here, we analyzed the functional profile of peDNA for each mode of transportation, EV- and GTA-mediated gene transfer, and transduction. In brief, each peDNA read that mapped to either an EV-, GTA producer or a transducer was classified into a cluster of orthologous groups (COG), and the resulting profile was then normalized with the profile of microbial reads (corresponding metagenome) mapping the respective sample (Figure 7). The COG category ‘Mobilome (Prophages, transposons)’ was overrepresented in all three groups. For transducers, this overrepresentation is mainly due to an actively transducing provirus present in the peDNA. However, the overrepresentation of the COG category ‘Mobilome’ in GTA and EV derived peDNA, is significant and we hypothesize, this is due to the presence of transposons. In fact, 82 % of all reads assigned to the category ‘Mobilome’ were assigned to a cluster with the keyword ‘transposon’. We suggest that transposon activity is also the reason for the overrepresentation of the other COG categories: ‘Extracellular structures’, ‘Signal transduction mechanisms’, and ‘Cell motility’. All three categories are associated with the adaptation of the organism to a changing environment. Genes of these categories are often found on ‘genomic islands’ (GIs), highly variable and mobile regions on the genome (52, 53). At the same time, the occurrence of transposons on GIs is well-documented. Transposons have been shown to mobilize not only themselves but also adjacent ’passenger genes’, genes that are located in proximity to transposons and are therefore co-mobilized by transposons (54). Evidence shows that environmental stressors increase the activity of transposons (55). as well as the production of EVs (56), and the induction of proviruses (57) and GTAs (27). Increased transposon activity increases the intracellular mobilization of genes surrounding transposons and therefore could lead to an increased uptake into EVs, GTAs or virus particles. Our data suggest that these two stress-induced mobilization mechanisms may be linked in a way that enhances the community’s adaptability to the environment, by increasing genetic transfer between individual cells.

**Figure 7:**
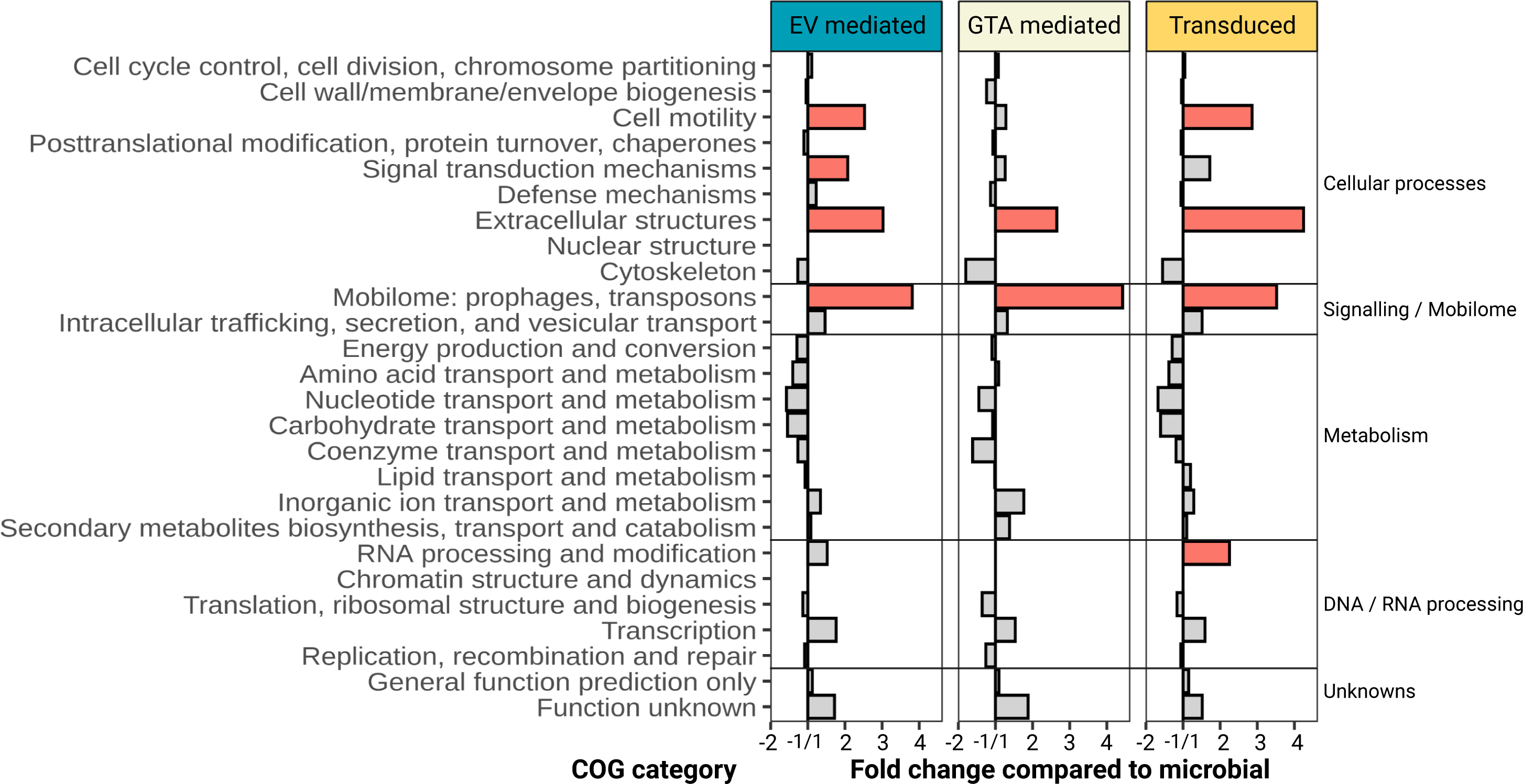
Frequency of overrepresentation of clusters of orthologous groups (COG) categories for peDNA assigned to EV- and GTA producer and Transducer. Bars represent the frequency of overrepresentation of genes assigned to each category, for each type of MAG (GTA producer, EV producer, Transducer). Red bars indicate that a higher percentage of genes belonging to this category showed increased (two times standard deviation above mean) recruitment rates of peDNA reads.

## Conclusion and Outlook

In this study, we propose the term ‘protected extracellular DNA’ (peDNA) to refer to genetic material obtained from size-filtered and DNase-treated samples, thereby accommodating non-viral, EV- or GTA-transported DNA in that sequence space. So far, there is no known sequence marker to distinguish horizontally transferred DNA from cellular DNA, therefore the removal of contaminating cells and free DNA is crucial when analyzing peDNA samples. The level of contamination however, should not be assessed using the presence of ribosomal subunit-mapping reads, since EVs have been shown to transport 16/18S rRNA genes.

In our study, we analyzed a carefully purified marine sample of peDNA and existing global marine datasets. We were able to link peDNA sequences to potential hosts and identify their primary mode of DNA transfer. Among the identified GTA and EV producers, most were shown to produce the respective particles in previous studies, confirming the validity of our approach, however, new potential GTA and EV producers were also identified. Overall, EV-mediated gene transfer was the most common mechanism and we hypothesize that EVs are a main driver of HGT in the ocean. Lastly, our findings suggest that EV-mediated gene transfer and transposon induced gene mobilization potentially work together and enhance the ability of microbial communities to adapt to a changing environment. Given the considerable ecological stressors imposed by climate change, comprehensive investigations into the role of EVs, GTAs and viruses for HGT is essential to understand genetic adaptability in marine microbes. This study highlights the need for further research into HGT mechanisms, and peDNA in general, since the community composition and function of marine microbes, and therefore the global oceans, is strongly shaped by the abundance of protected viral and non-viral extracellular DNA.

## Authors Contributions

D.L. performed the majority of the experimental laboratory and bioinformatic work. C.M. and T.A.S. supported general laboratory work, sampling and purification of sea water samples. S.E. conceived and led the study. D.L. and S.E. performed the primary writing of the manuscript. All authors participated in the analysis and interpretation of the data and contributed to the writing of the manuscript.

## Acknowledgements

We thank the sampling crew of the Heligoland sampling station ‘Kabeltonne’ for their help obtaining the sea water samples and the AWI research station on heligoland for providing laboratory workspace. We are grateful to Luis ‘Coto’ Orellana and Isabella Maria Wilkie for providing access to their data. Finally, we thank A. Probst, A. Zayed and B. Fuchs for insights, discussions and feedback. We thank Daniela Thies and Ingrid Kunze (MPI for Marine Microbiology, Bremen, Germany) for assistance with some of the experiments. Finally, we want to thank the Max-Planck-Institute for Marine Microbiology and the Max-Planck-Society for continuous support.

## Data Availability

Heligoland metagenome and peDNA reads are available at the European Nucleotide Archive (ENA) under BioProject PRJEB60526.

## Methods

### Sampling and Filtration

A visual overview of the sampling and filtration methods is given in Figure S6. Three seawater samples (G, H, I) of 100 liters were collected off the shore of Helgoland at the sampling station ‘Kabeltonne’ (54°11’02.4’N 7°53’49.2’E). Each sample was sequentially filtered through 10 µm, 3 µm, 0.8 µm, 0.45 µm and 0.22 µm (polyethersulfone filters, Merck Millipore, Burlington, MA, US). Filters were immediately stored at -20° C for later DNA extraction. Flow through of the 0.22 µm filters was subsequently concentrated using tangential flow filtration with a 100 kDa cassette (Sartorius Stedim). The concentrated samples were stored at 4 °C, until further concentration down to 0.5 ml, using Amicon filter centrifugation (1 MDa AmiCon tube filters, 2500 x g). Finally, the concentrated sample was diluted with purified seawater (flow-through from the tangential flow filtration ) to 2 ml. Two aliquots were created for each sample á 0.5 ml, one treated with DNase before gradient purification, one treated afterwards (see ’Purification of peDNA samples’).

### Purification of peDNA samples

In order to remove free DNA, half of the samples were incubated with 100 U / ml DNase I at 37°C for 10 minutes, while the other half were DNase-reated after density gradient purification. EDTA was added to a final concentration of 5 mM and the enzyme was deactivated at 75°C for 10 minutes. CsCl density gradients were prepared as following: Five CsCl solutions were prepared with 25, 30, 35, 40 and 60 % CsCl solved in artificial sea water (480 mM NaCl, 27 mM MgCl2, 2.8 mM MgSO4, 9 mM KCl, 6 mM NaHCO3, 10 mM CaCl2) (58). For each gradient, 1 ml of each solution was carefully layered on top of each other and stored at 4 °C overnight in order to establish the gradient. 0.5 ml of sample was carefully placed on top of the samples, before ultracentrifugation (20 h, 38,000 x *g*, 4° C). For each gradient, individual 0.5 ml fractions were carefully extracted and incubated with 40 % PEG 6000 (final concentration 10 %) overnight at 4 °C. Particles were precipitated by centrifugation (13,000 x *g,* 45 min, 4 °C) and particle pellets dissolved in 1 ml artificial sea water. The other half of the samples were DNase-reated at this time point. A detailed overview of the samples is given in Table S1 and a visual overview in Figure S7.

### DNA Extraction

Frozen polycarbonate filters (3, 0.8, 0.45 and 0.22 µm) were placed in a 50 ml tube together with 13.5 ml of extraction buffer (100 mM Tris-HCI pH 8.0, 100 mM EDTA pH 8.0, 100 mM Na-Phosphate buffer pH 8.0, 1.5 M NaCI, 1 % CTAB). peDNA samples were processed directly. DNA was extracted as described elsewhere (59). In brief, samples were treated with 10 mg/ml Proteinase K and incubated at 37 °C for 30 min on a shaker. Then, 1/10 vol of 20% SDS was added, before incubating again at 65 °C for 2 h on a shaker. After centrifugation (53,000 x *g*, 10 min, RT), the samples were transferred into a new tube and 1 vol of chloroform/isoamylalcohol was added and samples were thoroughly mixed, before centrifuging at 4000 x*g* for 20 min at RT. The aqueous phase upper phase was collected and transferred into a new tube. This step was repeated until no protein/polysaccharide layer was visible. DNA was then precipitated by adding 0.6 vol isopropanol and incubation for 1 h at room temperature. DNA was pelleted at 53,000 x *g* for 10 min at RT washed with 1 ml cold (4 °C) 80 % ethanol and resuspended in 60 µl 1x TE buffer overnight. DNA concentration was assessed using a spectrophotometer (DS-11 FX + by DeNovix**®**, Wilmington, DE, US), see Table S1.

### Sequencing

DNA samples were pooled according to Table S1, assuring enough DNA content per sample for successful sequencing. Library preparation (FS DNA Library, NEBNext® Ultra™, Ipswitch, MA, US) and sequencing (Illumina HiSeq2500 by Illumina, San Diego, CA, US, 2 x 250 bp for peDNA samples and Illumina HiSeq3000, 2 x 150 bp for Filter DNA) was performed at the Max Planck Genome Centre Cologne (MP-GC).

### Read trimming and assembly

Paired-end reads from Heligoland EV enrichments were trimmed using Trimmomatic [Bolger 2014] in paired-end mode, with the parameters LEADING:8 TRAILING:8 SLIDINGWINDOW:5:24 MINLEN:50. Paired-end reads from EV Enrichments (33) and paired-end reads from GDOCB (42) were trimmed using bbduk.sh, part of the BBTools suite (60) with the following parameters: <u>bbduk.sh</u> qtrim=rl trimq=20 maq=20 minlen=30 ordered t=8 ref=adapters.fa, where adapters.fa were fasta files containing adapters identified to be present in the reads using FastQC (61). Reads from Heligoland EV enrichments were assembled using metaSPAdes (62) with default parameters.

### Handling of external data

External datasets were downloaded from public servers. An overview of external datasets used in this study, with SRR, ERS and DRR accessions, is given in Table S1. Reads from Tara Ocean viromes (17) were already trimmed. Reads from EV enrichments (33) and GDOCB viromes (42) were assembled using metaSPAdes (62) with default parameters. For Tara Oceans viromes, assembled contigs were downloaded from https://www.ebi.ac.uk/. Tara Oceans MAGs were published elsewhere by Tully et al (44), accessions are listed in Table S1.

### Calculation of SSU alignment rates

SSU alignment rates were calculated using ViromeQC (19), which maps input reads against 16S and 16S rRNA subunits. This was done for all Tara Ocean viromes, Heligoland peDNA, EV enrichments (see Table S1 for an overview of all samples used).

### Calculation of the percentage of non-viral associated reads

The percentage of non-viral peDNA in viromic samples was calculated with a pipeline of bioinformatic tools. An overview is given in Figure S4. Paired-end, trimmed input reads were assembled. Contigs shorter than 2000 bp were removed from downstream analysis. Then, contigs were subject to two viral-prediction steps: viral sequences were predicted i) using a combination of VirSorter2 and CheckV as described previously (63) and ii) DeepVirFinder (16). The results of both steps were summarized using custom script and each contig was labeled as either ’viral’ or ’non-viral’. Then, the initial input reads were mapped against labeled contigs, using bbmap.sh, part of the BBTools suite (60) with default parameters. Then the number of non-viral-contig mapping reads was divided by the number of total reads mapping against viral or non-viral contigs. This ratio of non-viral/viral reads is referred to as ’percentage of non-viral to viral associated reads’ or ’nvpeDNA / peDNA read ratio’ in this study.

### Identification of potential transducers, extracellular vesicle- and gene transfer agent producers

In order to identify potential EV producers, GTA producers and MAGs with an actively transducing virus, a second bioinformatic pipeline was developed (see Figure S5). First, MAGs were filtered by removal of MAGs shorter than 100,000 bp. Then, reads from the corresponding viromes / peDNA samples were mapped against the MAGs, using bbmap.sh with default parameters. For each sample (in total 9 samples from Tara Oceans, 1 combined Heligoland sample) the twenty most recruiting MAGs were selected for further downstream work. VirSorter2 (default parameters) was used in order to predict potential integrated proviruses. GTA clusters were predicted by searching for homologs of proteins of known GTA clusters using *diamond blastp* with default parameters (evalue <= 10^-5, pident > 50 %). Finally, a custom in-house script summarized the results and an automated label was given. Additionally, each label was manually curated and each MAG was labeled as either EV producer, GTA producer or an organism with an actively transducing virus (see Figure S5 and S6).

### Annotation of GTA producers and viral regions

Open reading frames were predicted using prodigal with the metagenome flag (*prodigal -i <FASTA-file> -d <GENES-out> -a <PROTEIN-out> -p meta*). Each ORF was then annotated using the InterProScan API (https://github.com/ebi-wp/webservice-clients-generator) with default parameters (64) and additionally checked manually. For the prophage region, shown in Figure 6, the DNA sequence was submitted and annotated in PHASTER (65, 66). Genome maps and presence-absence plots were generated using ggplot (67) and BioRender.com.

### Identification of Cluster of orthologous groups

In order to assess the functional profile of peDNA reads, each read was mapped to the 170 MAGs for which the primary transport mechanism was identified using bbmap, part of the BBTools suite (60) with minid=95 and otherwise default parameters. This resulted in three sets of reads: EV-mediated, GTA-mediated, VLP-mediated. For each read partial ORFs were predicted using FragGeneScan (68) with the parameters -complete=0, -train=illumina_5 and otherwise default parameters. The partial ORFs were then blasted against the COG database (69) using the diamond tool set (70) with the following parameters: *-f 6 –max-target-seqs 1 – query-cover 80 –subject-cover 10*. Each read was then assigned a COG cluster and consequently a COG category. For each category, the relative abundance was calculated using

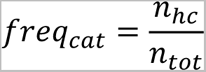

where *n_hc* is the number of reads in category *cat* that show high coverage and *n_tot* is the total number of reads assigned by this label. The same procedure was done for metagenome reads of the corresponding metagenome samples (see Supplementary Table S1 - Sample Overview). The fold change between categories was calculated pairwise with the formula

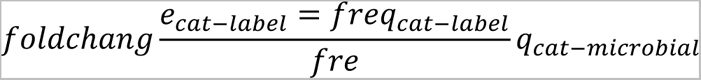

where *cat* refers to a specific COG category, label to either EV, GTA or virally transduced and microbial to the microbial counterpart of that sample. For visualization reasons, fold changes smaller 1 were calculated with the reversed formula

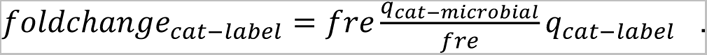

Fold changes between -1 and 1 are therefore not possible and this area is excluded from the plot.

### Coverage plots, genome maps and schematic figures

Coverage plots of potential transducers, EV- and GTA producers were created using the R package ggplot2 (67). Genome maps of potential GTA producers were created using the R package gggenes (https://github.com/wilkox/gggenes). Schematic genome maps and additional elements in figures were created with BioRender.com.

