## Supplemental Material for "Extracellular vesicles are the main contributor to the non-viral protected extracellular sequence space"

### Supplementary Material


**Supplementary Material 1** **1**

Figure S1: Pipeline 1 - Calculation of “non-viral / viral read ratio” or “percentage of non-viral associated reads”. 2

Figure S2: Pipeline 2 - Identification of potential EV-, GTA producers and microbes with actively transducing virus. 3

Figure S3: MAG abundance in metagenomes versus virome/EV-enrichment. 3

Figure S4: Decision making logic for the identification of potential EV-, GTA producers and microbes with actively transducing virus. 4

Figure S5: Coverage plots (separate pdf file)

Figure S6: Overview of the sampling and purification workflow. 5

Figure S7: Overview of sequencing effort of CsCl-gradients. 7

Table S1: Overview of sheets in Table S1 (separate xlsx file) 7

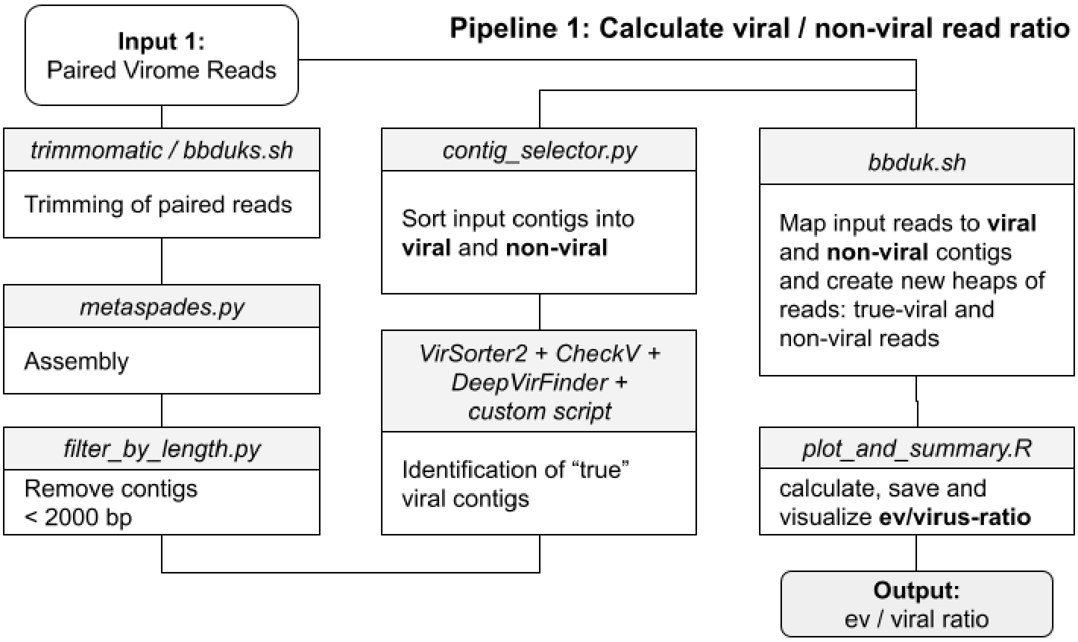


**Figure S1: Pipeline 1 - Calculation of "non-viral / viral read ratio" or "percentage of non-viral associated reads".**

Schematic overview of the bioinformatic pipeline in order to calculate the percentage of non-viral associated reads within a given virome / EV enrichment. Reads were trimmed, assembled into contigs and short contigs removed. Each contig was labeled as "viral" or "non-viral" based on the results of virus prediction tools. Finally, the input reads were mapped against the contigs and the ratio between viral-mapping and non-viral-mapping reads was calculated.

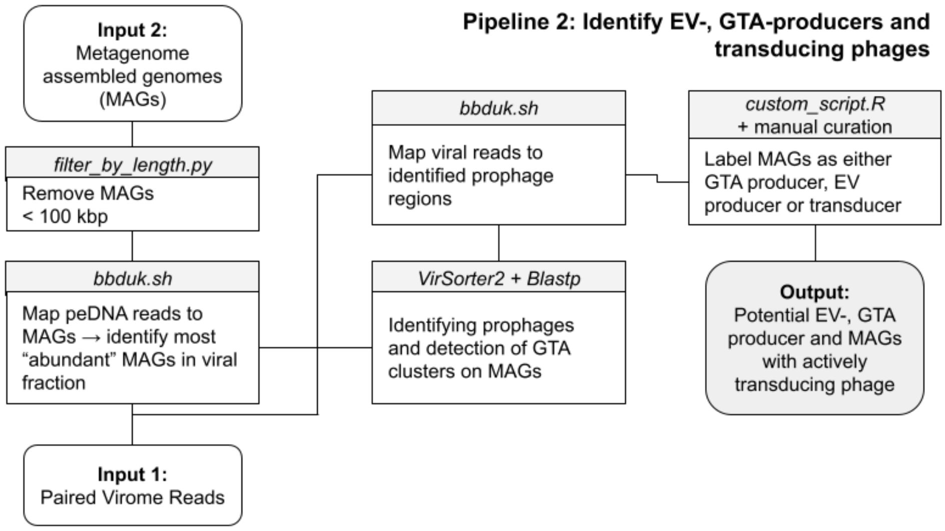


**Figure S2: Pipeline 2 - Identification of potential EV-, GTA producers and microbes with actively transducing virus.**

MAGs shorter than 100 kbp were removed. peDNA reads were mapped against each MAG and the 20 most peDNA-recruiting MAGs were selected for further analysis. Each MAG was subsequently scrutinized for the presence of a prophage or a GTA cluster and subsequently labeled as either GTA producer, EV producer or transducer using an in-house custom script and manual curation (decision making logic, see Figure S6).

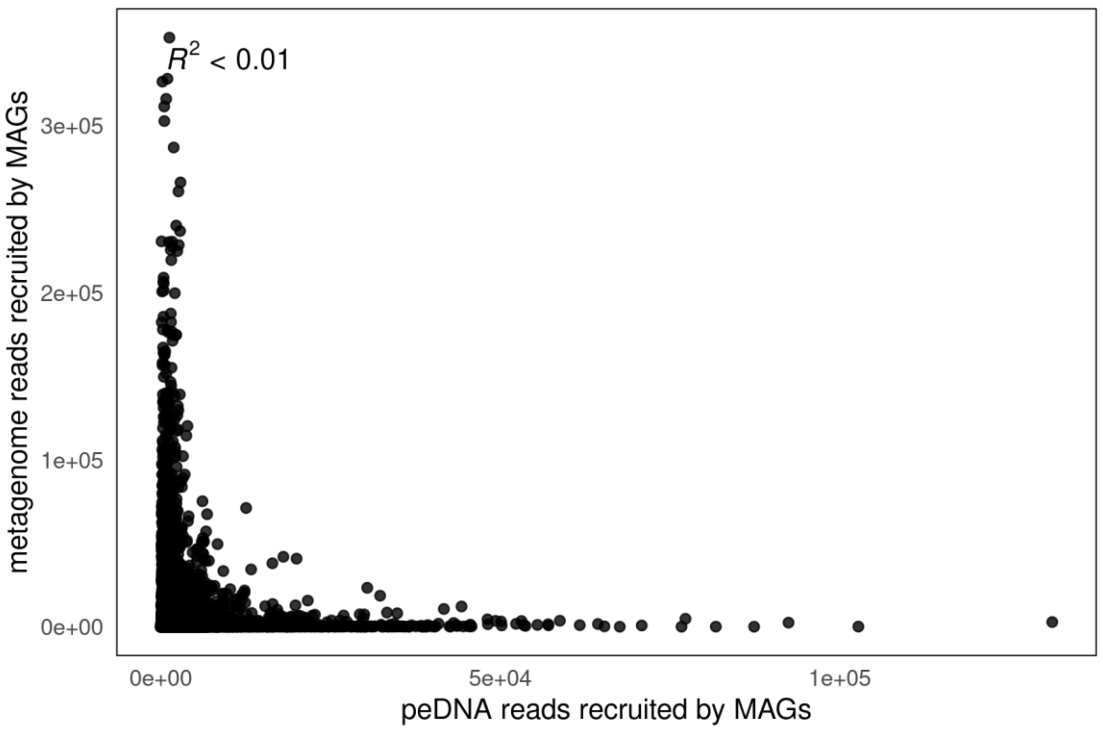


**Figure S3: MAG abundance in metagenomes versus virome/EV-enrichment.**

Number of virome reads recruited on the x-axis, number of metagenome reads recruited on the y-axis, per MAG. R^2 = 0.003391 calculated using linear regression.

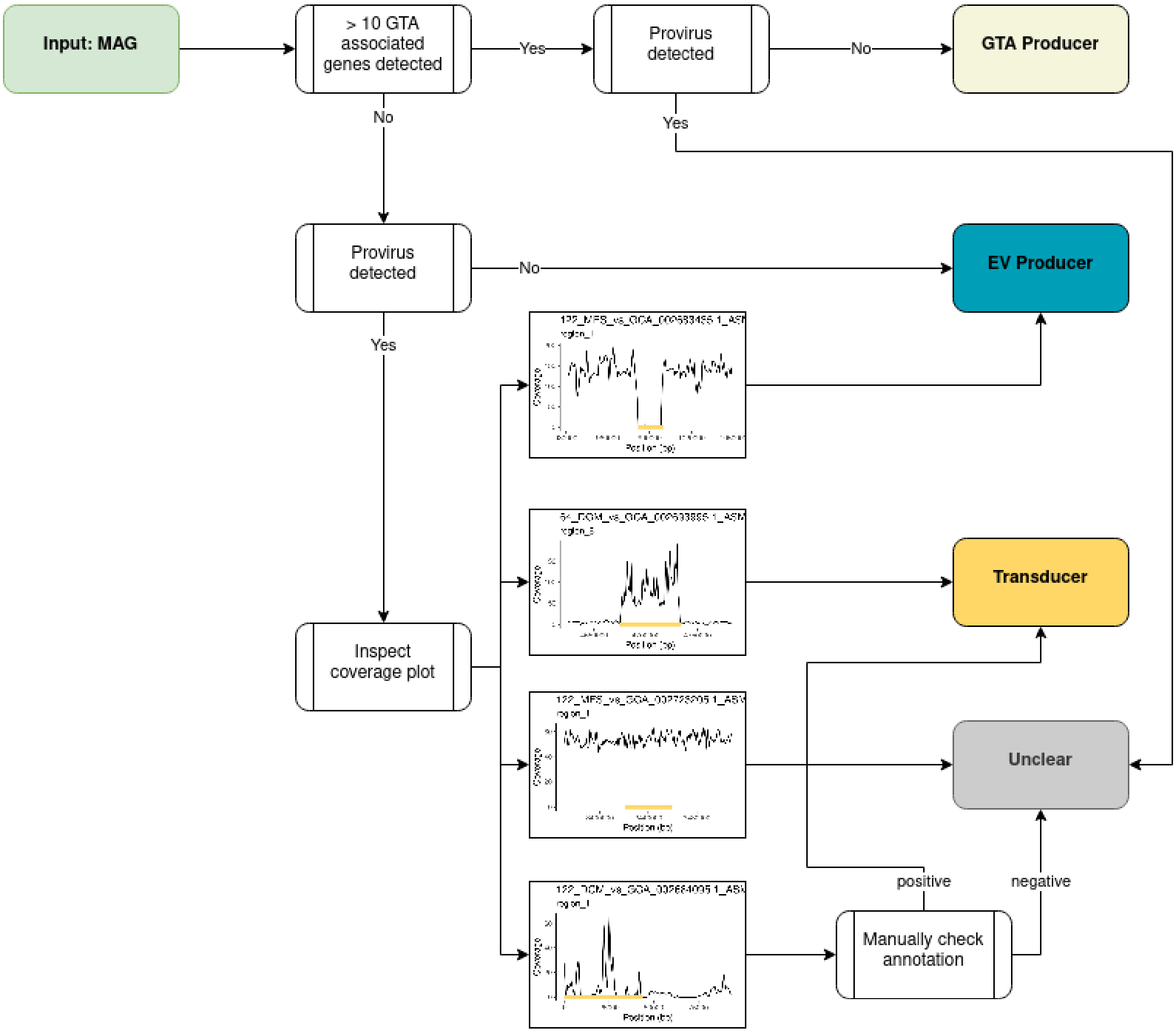


**Figure S4: Decision making logic for the identification of potential EV-, GTA producers and microbes with actively transducing virus.**

A MAG was labeled as “GTA producer” if >10 GTA associated genes were detected and no provirus was found. If <10 GTA genes and no provirus was found, the MAG was labeled “EV producer”. If both a provirus was detected and >10 GTA genes, the MAG was labeled “unclear”. If a provirus was detected, the coverage plot was scrutinized for manual inspection and decision making. If the viral region was “absent”, the MAG was labeled “EV producer”. If the viral region showed an increased coverage above the region, it was labeled as “Transducer”. For unclear cases, the exact annotation of the viral region was analyzed and manually curated and combined with the coverage plot. Based on this information, the MAG was labeled either as “unclear” or “Transducer”.

See file: Figure S5 - Coverage Plots

**Figure S5: Coverage plots (Separate pdf file)**

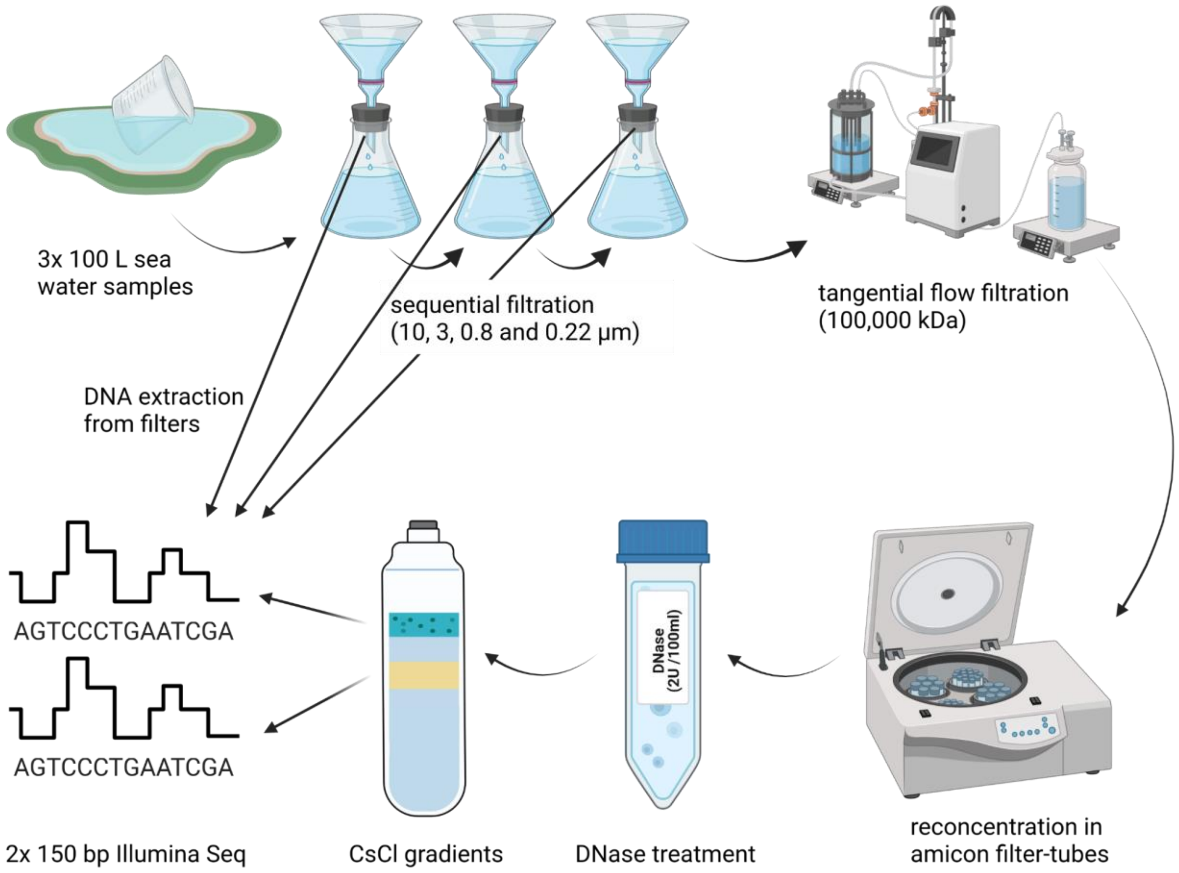


**Figure S6: Overview of the sampling and purification workflow.**

For details refer to the section “Methods - Sampling and Filtration”. In brief: Samples were sequentially filtered, concentrated using tangential flow filtration, further concentrated using amicon filter tubes, DNase treated, gradient purified using CsCl gradients. DNA from bands/fractions was extracted and sequenced.

DNA from the 3 µm, 0.8 µm and 0.22 µm filters was extracted and sequenced as well.

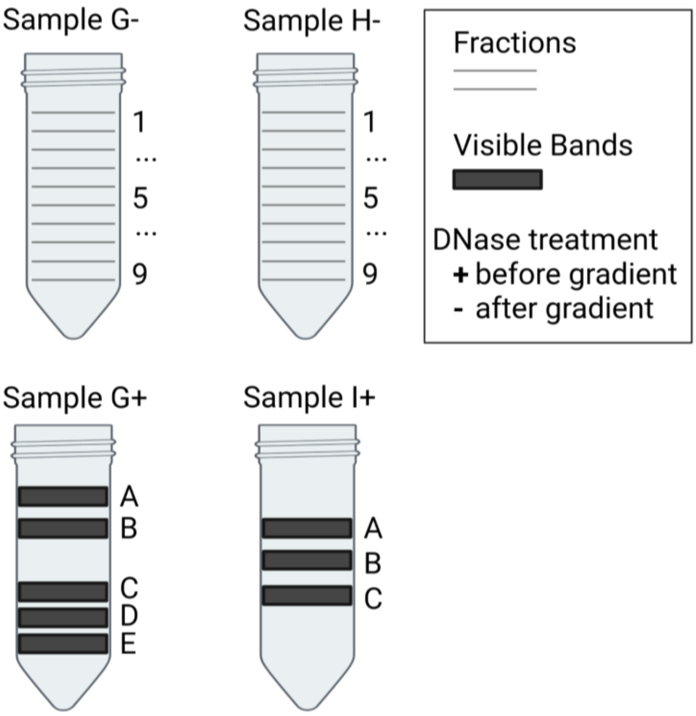


**Figure S7: Overview of sequencing effort of CsCl-gradients.**

Schematic overview of the samples used in this study. Samples marked with a + are treated with DNase, samples marked with a - are untreated. Sample names G, H and I refer to 3 different biological replicates (see Sampling and Filtration chapter). If bands were visible, bands were extracted and sequenced. If none were visible, fractions of 0.5 ml were extracted and sequenced.

**Table S1:** File - Table S1.xlsx

Overview of sheets given in Tabls S1:

| **Sheet** | **Description** |
|---|---|
| "Sample Overview" | Overview of all sea water samples analyzed. Information on sample processing and sequencing result |
| "peDNA/virome Datasets" | Run/Sample accessions for all 4 external peDNA/virome datasets used in this study |
| "Tully MAGs" | Overview of MAGs used in this study coming from Tully et al 2018 |
| "Orellana MAGs" | Overview of MAGs used in this study coming from Orellana et al 2019 |
| "top200 MAGs" | Summary of the top 200 peDNA-recruiting MAGs |
