## Supplementary material for "Extracellular vesicles are the main contributor to the non-viral protected extracellular sequence space": Figure S5

### Figure S5 - Coverage Plots

Binned (bin size = 1000 bp) coverage plots of all analyzed MAGs.

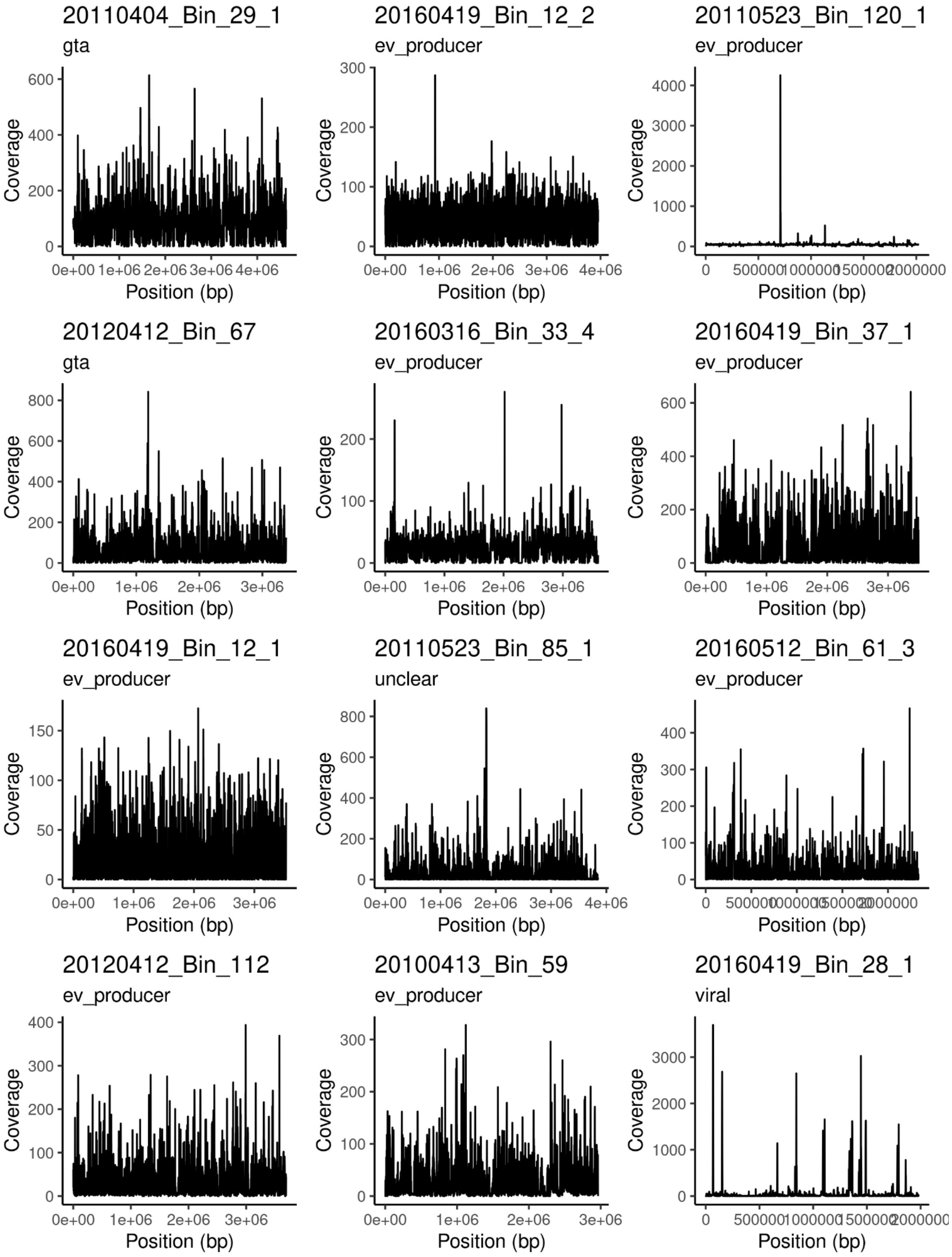
20110404_Bin_29_1
gta
20160419_Bin_12_2
ev_producer
20110523_Bin_120_1
ev_producer
20120412_Bin_67
gta
20160316_Bin_33_4
ev_producer
20160419_Bin_37_1
ev_producer
20160419_Bin_12_1
ev_producer
20110523_Bin_85_1
unclear
20160512_Bin_61_3
ev_producer
20120412_Bin_112
ev_producer
20100413_Bin_59
ev_producer
20160419_Bin_28_1
viral
Coverage
Position (bp)

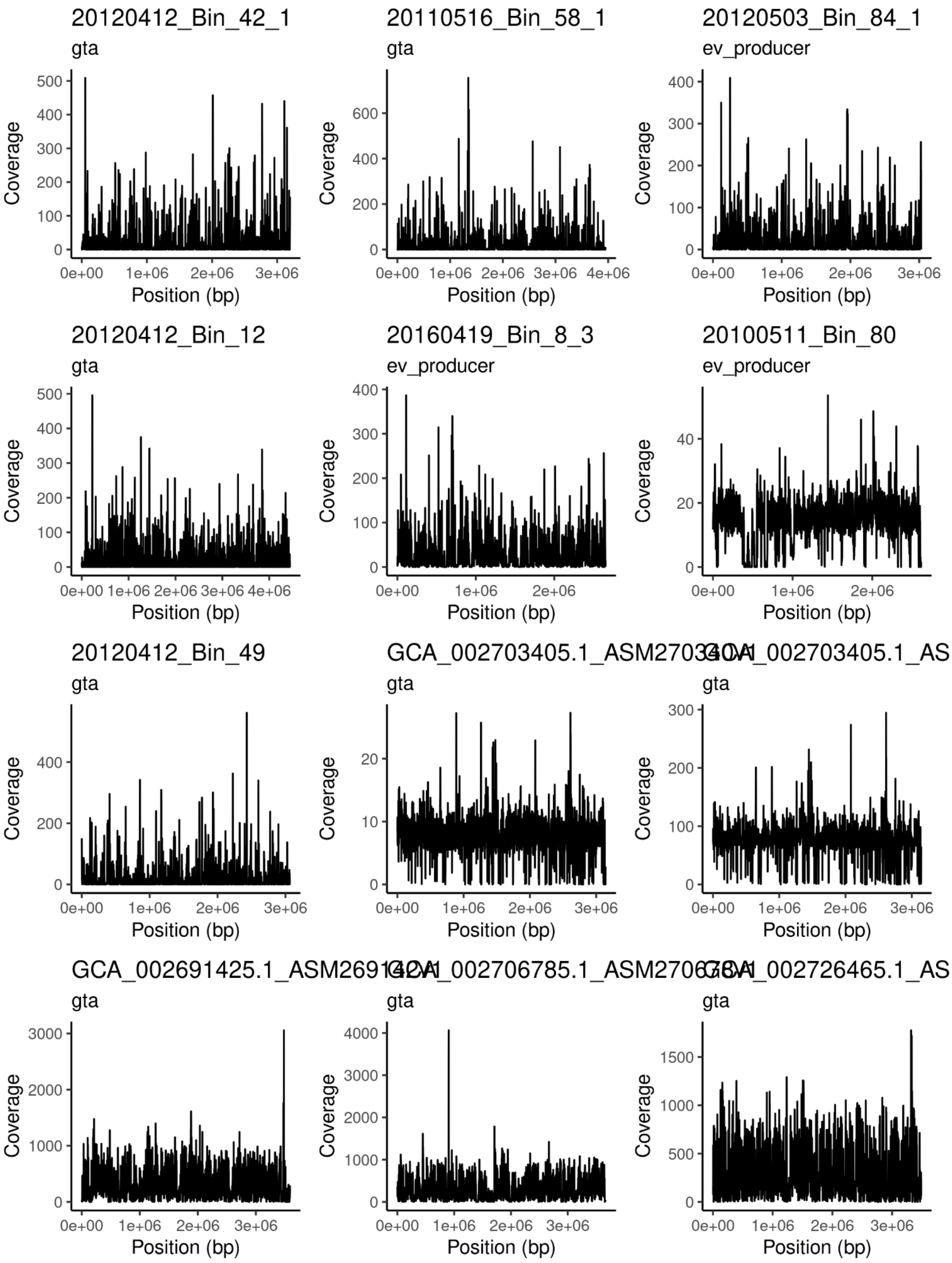
20120412_Bin_42_1
gta
20110516_Bin_58_1
gta
20120503_Bin_84_1
ev_producer
20120412_Bin_12
gta
20160419_Bin_8_3
ev_producer
20100511_Bin_80
ev_producer
20120412_Bin_49
gta
GCA_002703405.1_ASM27034
gta
GCA_002703405.1_AS
gta
GCA_002691425.1_ASM26914
gta
GCA_002706785.1_ASM27067
gta
GCA_002726465.1_AS
gta
Coverage
Position (bp)
0e+00
1e+06
2e+06
3e+06
4e+06

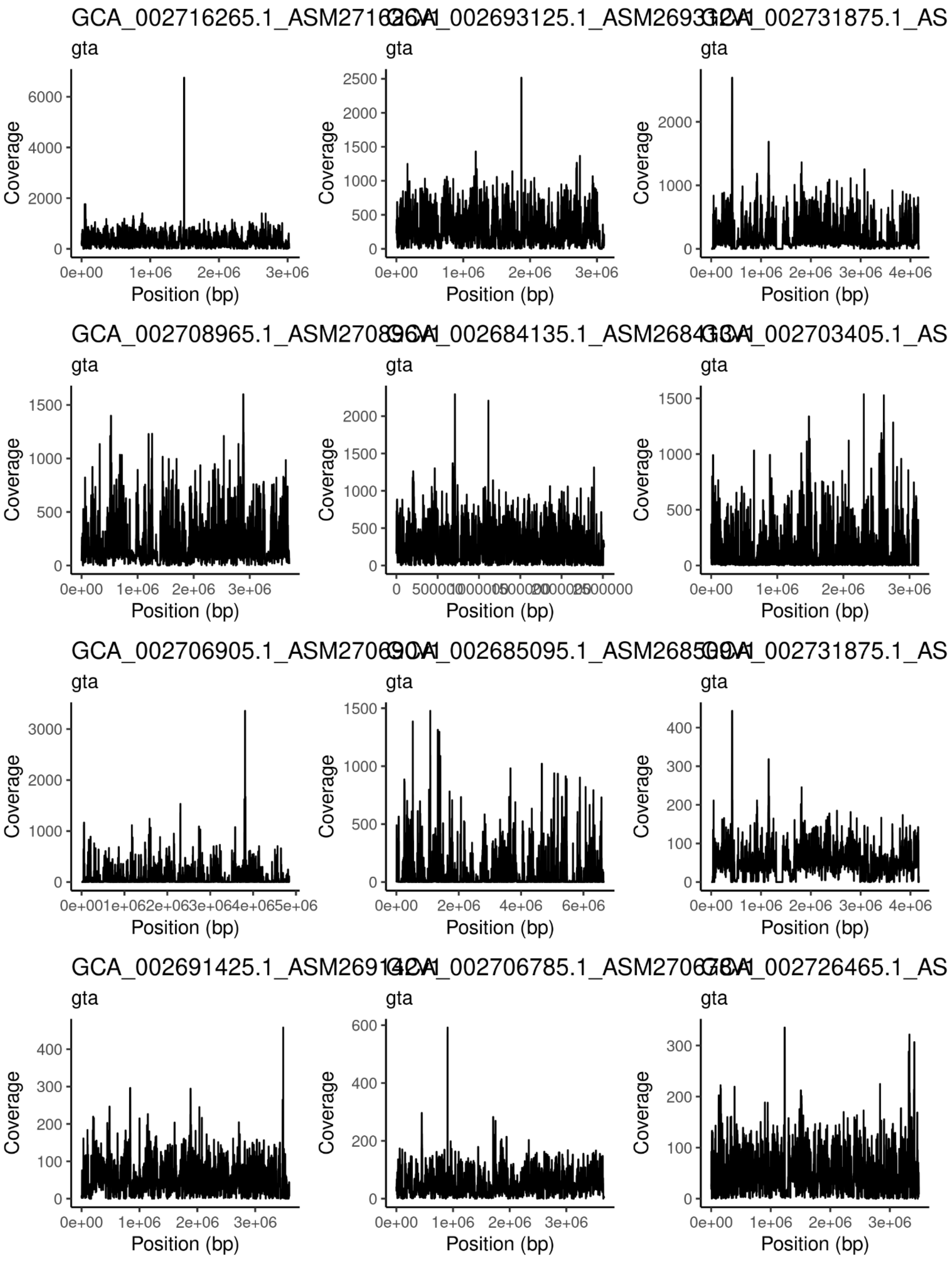
GCA_002716265.1_ASM2716
_002693125.1_ASM2693
_002731875.1_AS
GCA_002708965.1_ASM2708
_002684135.1_ASM2684
_002703405.1_AS
GCA_002706905.1_ASM2706
_002685095.1_ASM2685
_002731875.1_AS
GCA_002691425.1_ASM2691
_002706785.1_ASM2706
_002726465.1_AS
gta
Coverage
Position (bp)

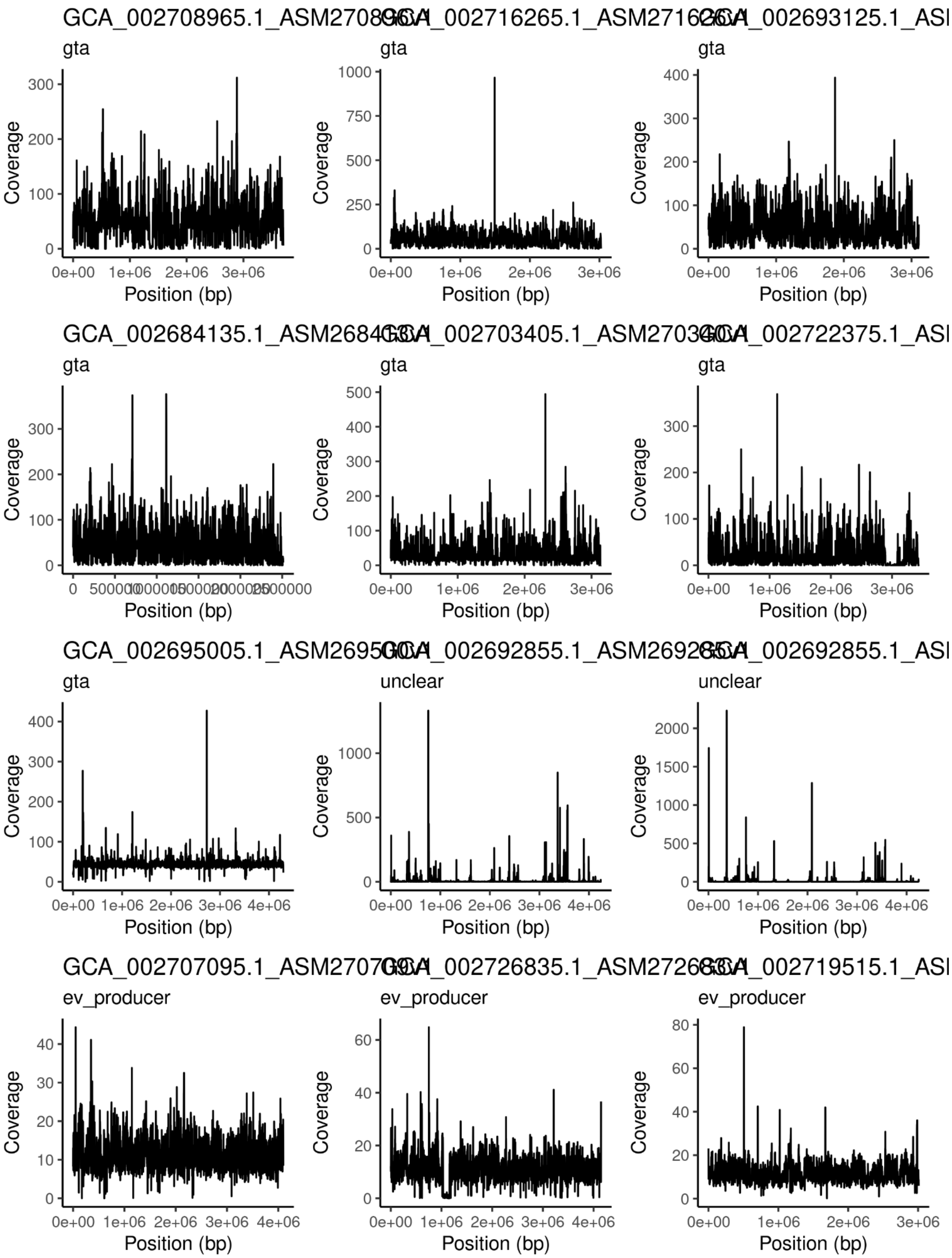
GCA_002708965.1_ASM27089
GCA_002716265.1_ASM27162
GCA_002693125.1_ASI
GCA_002684135.1_ASM26841
GCA_002703405.1_ASM27034
GCA_002722375.1_ASI
GCA_002695005.1_ASM26950
GCA_002692855.1_ASM26928
GCA_002692855.1_ASI
GCA_002707095.1_ASM27070
GCA_002726835.1_ASM27268
GCA_002719515.1_ASI
gta
unclear
ev_producer
Coverage
Position (bp)

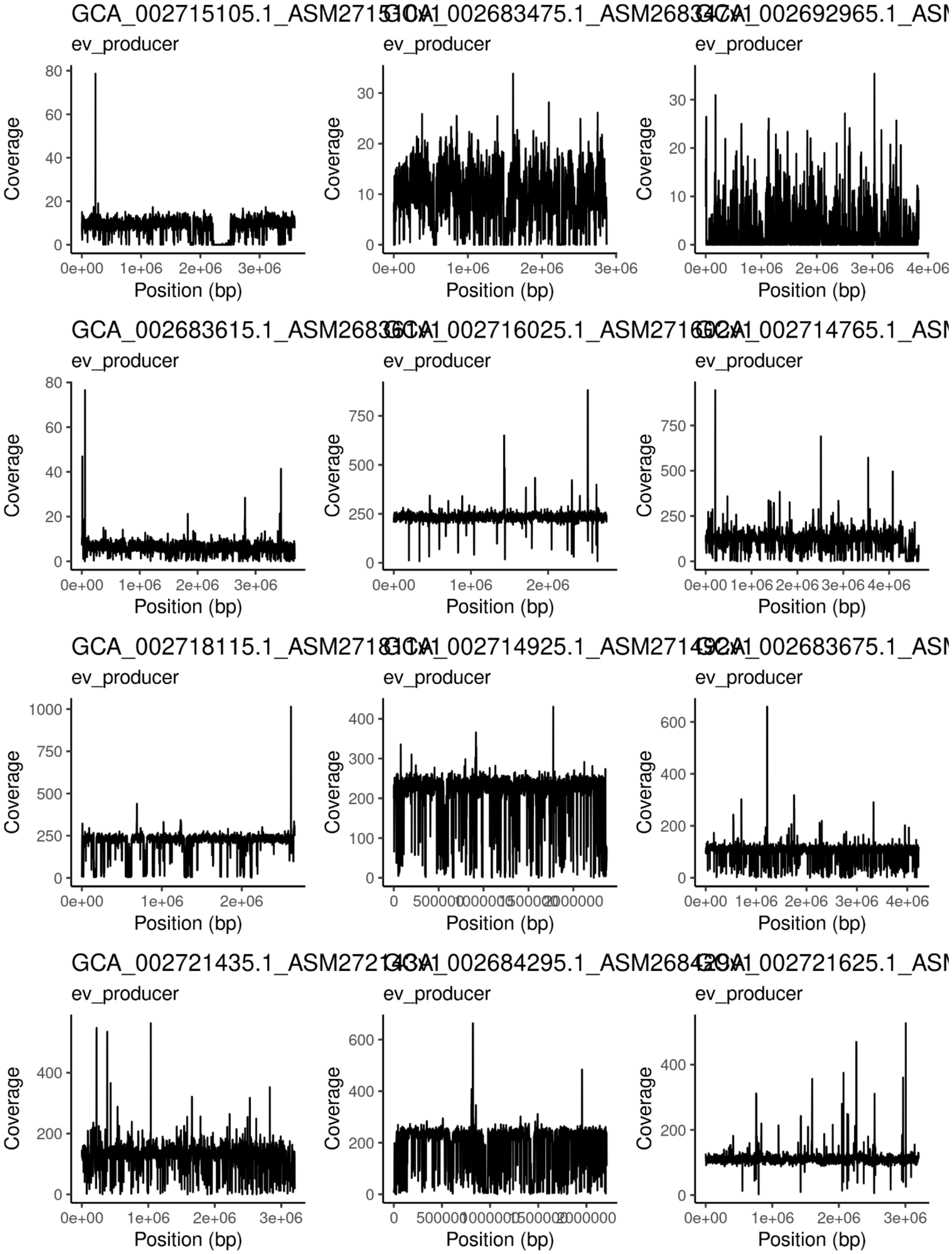
ev_producer
Coverage
Position (bp)

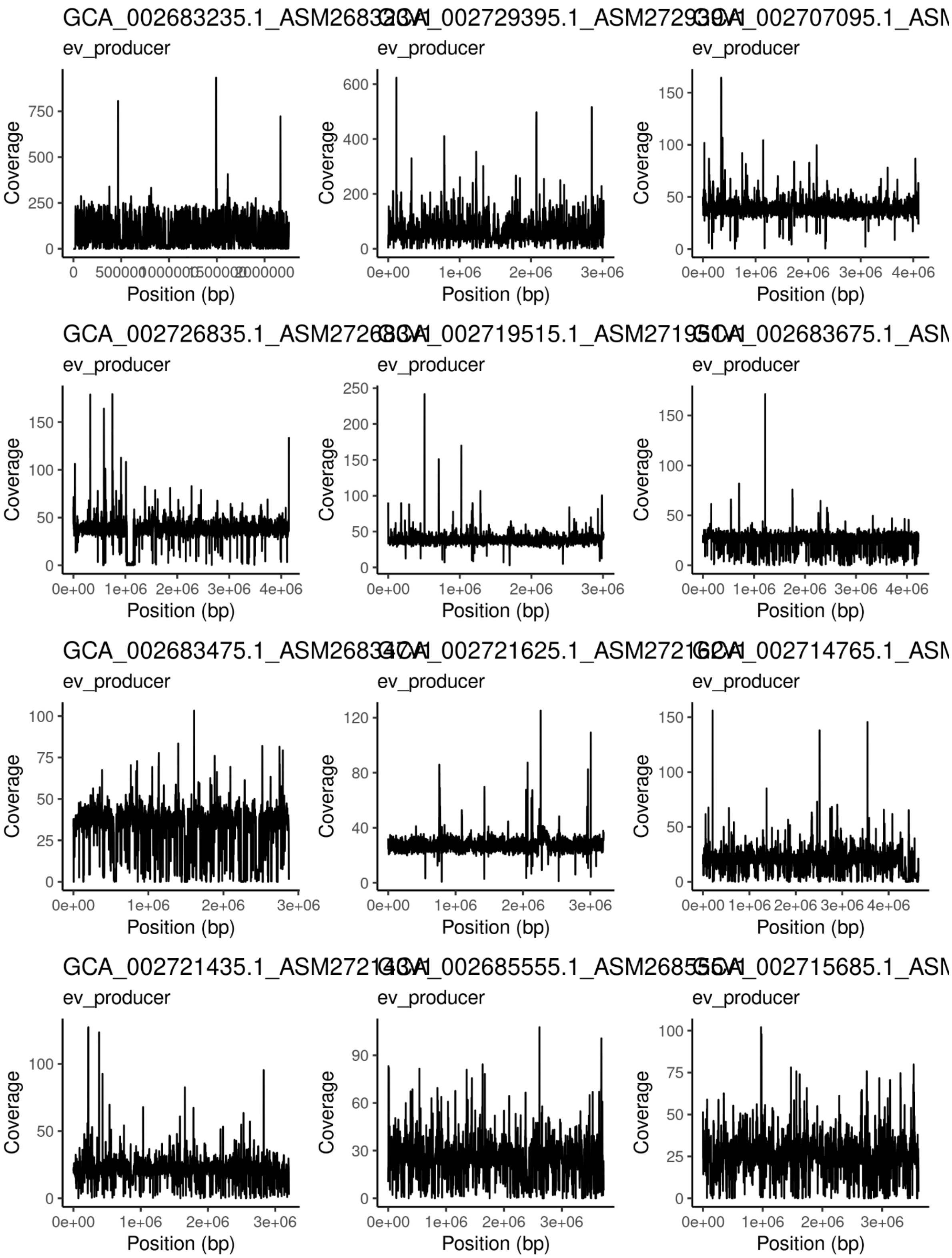
GCA_002683235.1_ASM2683
GCA_002729395.1_ASM2729
GCA_002707095.1_ASM
GCA_002726835.1_ASM2726
GCA_002719515.1_ASM2719
GCA_002683675.1_ASM
GCA_002683475.1_ASM2683
GCA_002721625.1_ASM2721
GCA_002714765.1_ASM
GCA_002721435.1_ASM2721
GCA_002685555.1_ASM2685
GCA_002715685.1_ASM
ev_producer
Coverage
Position (bp)

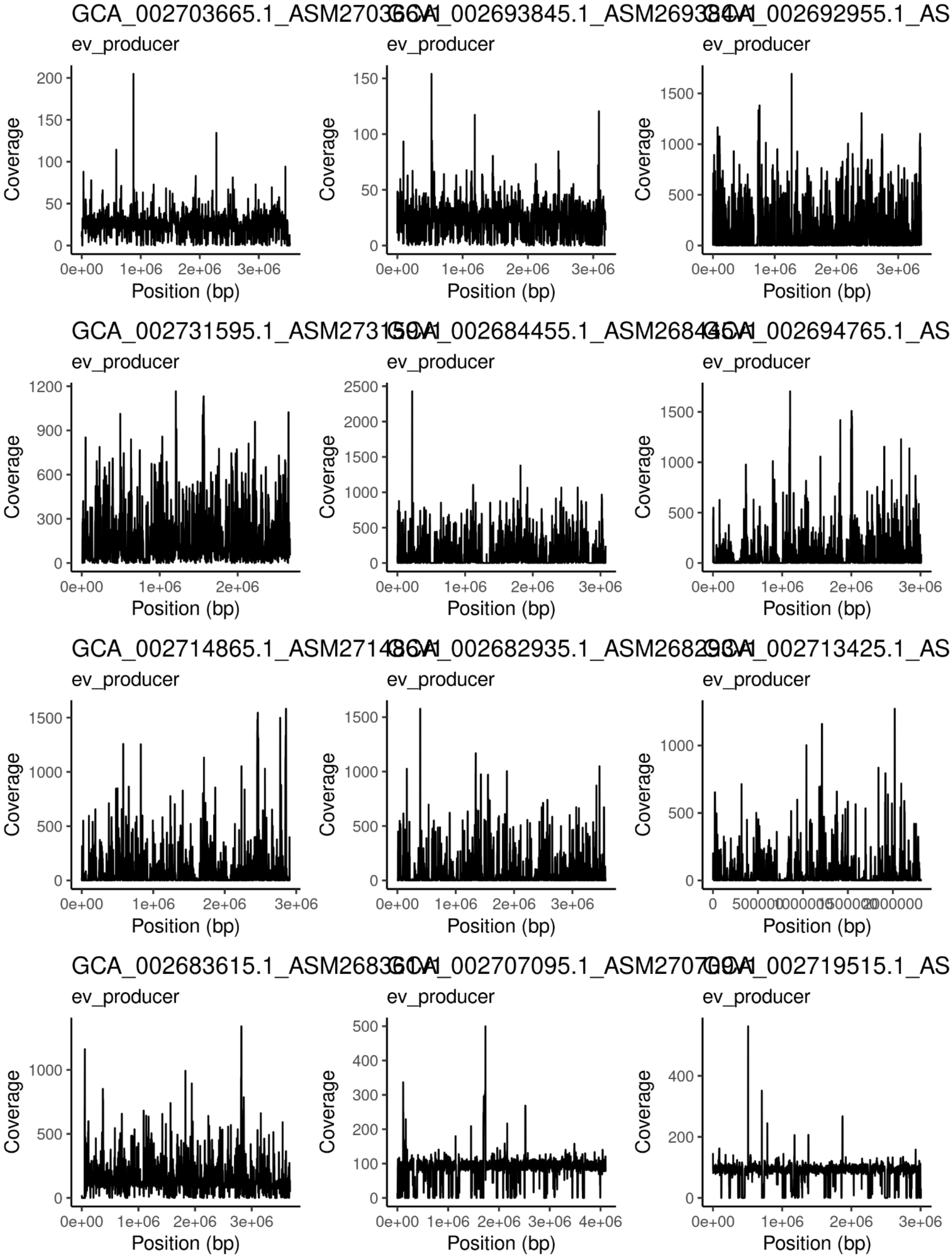

GCA_002703665.1_ASM270
GCA_002693845.1_ASM26938
GCA_002692955.1_AS
GCA_002731595.1_ASM2731
GCA_002684455.1_ASM26844
GCA_002694765.1_AS
GCA_002714865.1_ASM27148
GCA_002682935.1_ASM26829
GCA_002713425.1_AS
GCA_002683615.1_ASM26836
GCA_002707095.1_ASM27070
GCA_002719515.1_AS
ev_producer
Coverage
Position (bp)
0e+00
1e+06
2e+06
3e+06
4e+06
0
500000
2000000

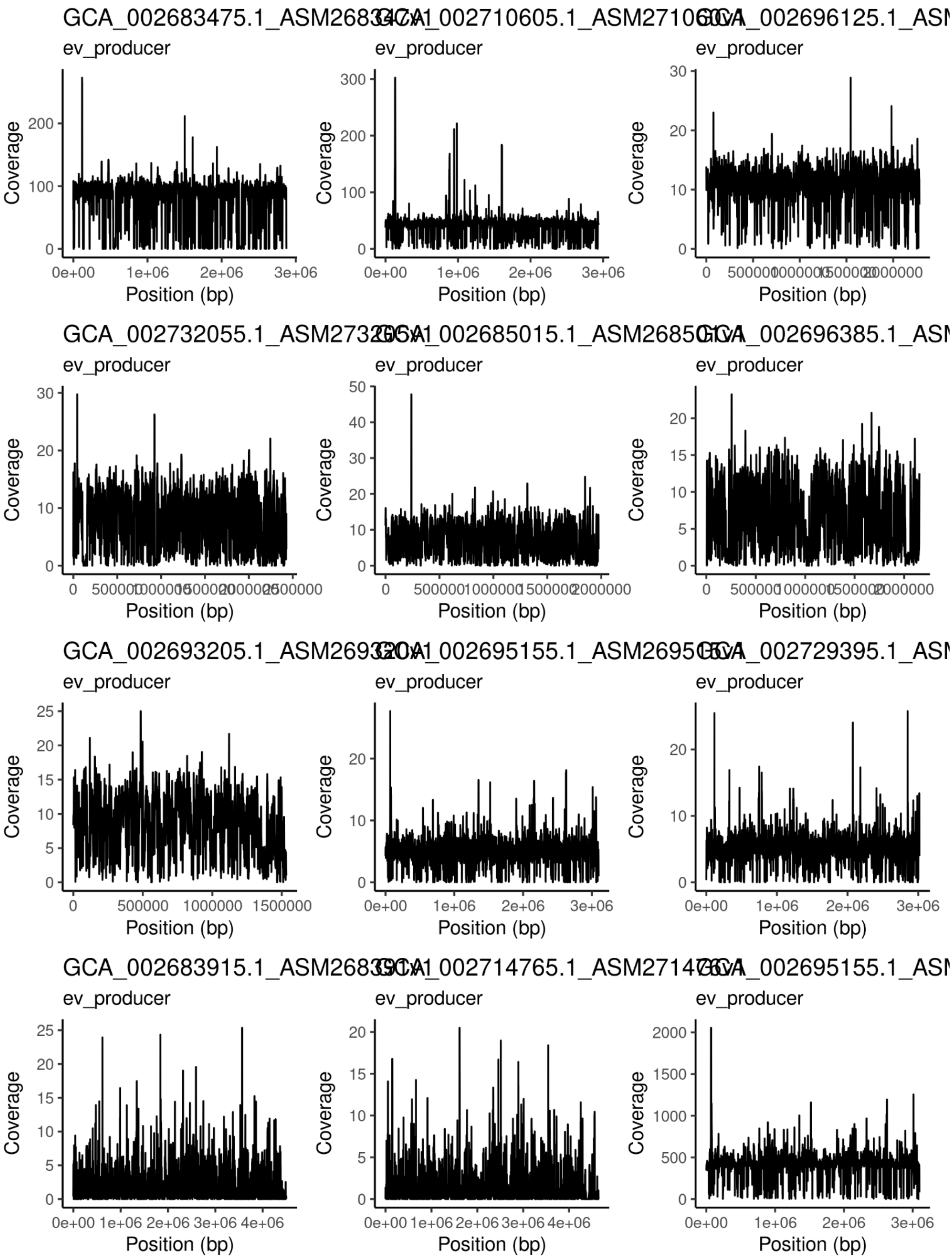
GCA_002683475.1_ASM2683
GCA_002710605.1_ASM27106
GCA_002696125.1_AS
GCA_002732055.1_ASM2732
GCA_002685015.1_ASM2685
GCA_002696385.1_AS
GCA_002693205.1_ASM2693
GCA_002695155.1_ASM2695
GCA_002729395.1_AS
GCA_002683915.1_ASM2683
GCA_002714765.1_ASM2714
GCA_002695155.1_AS
ev_producer
Coverage
Position (bp)

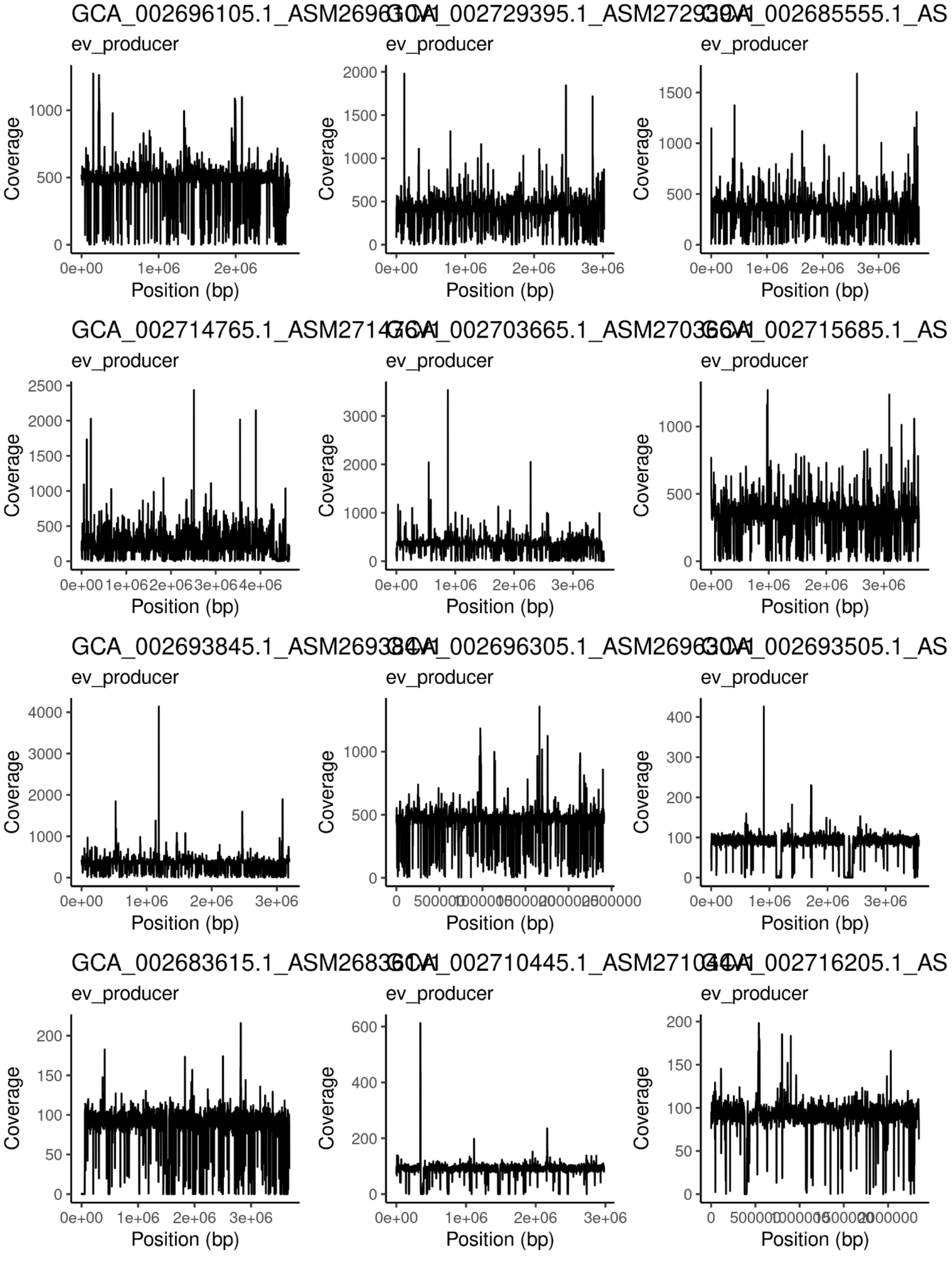
GCA_002696105.1
GCA_002729395.1
GCA_002685555.1_AS
GCA_002714765.1
GCA_002703665.1
GCA_002715685.1_AS
GCA_002693845.1
GCA_002696305.1
GCA_002693505.1_AS
GCA_002683615.1
GCA_002710445.1
GCA_002716205.1_AS
ev_producer
Coverage
Position (bp)

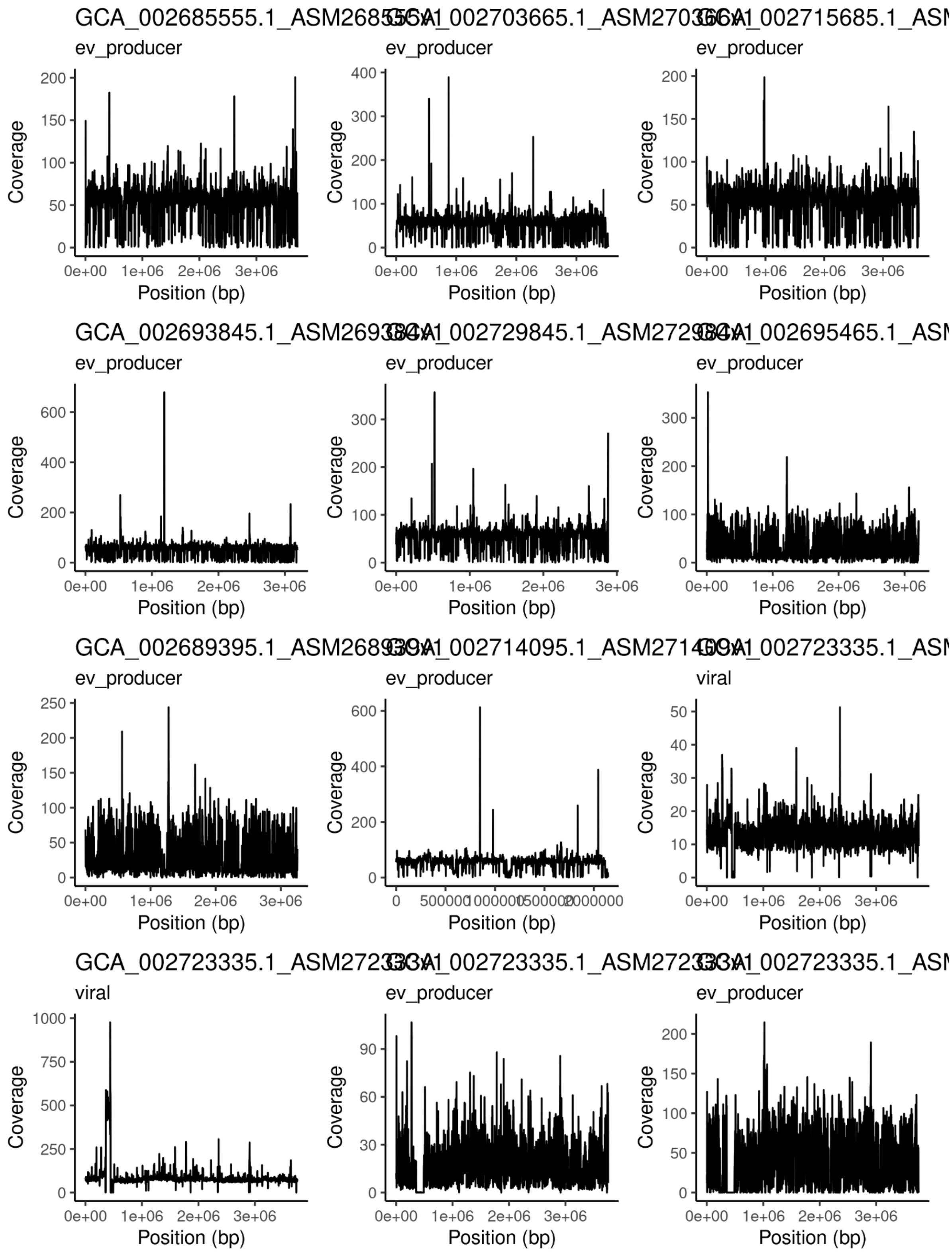
ev_producer
ev_producer
ev_producer
ev_producer
ev_producer
ev_producer
ev_producer
ev_producer
viral
viral
ev_producer
ev_producer
Coverage
Position (bp)

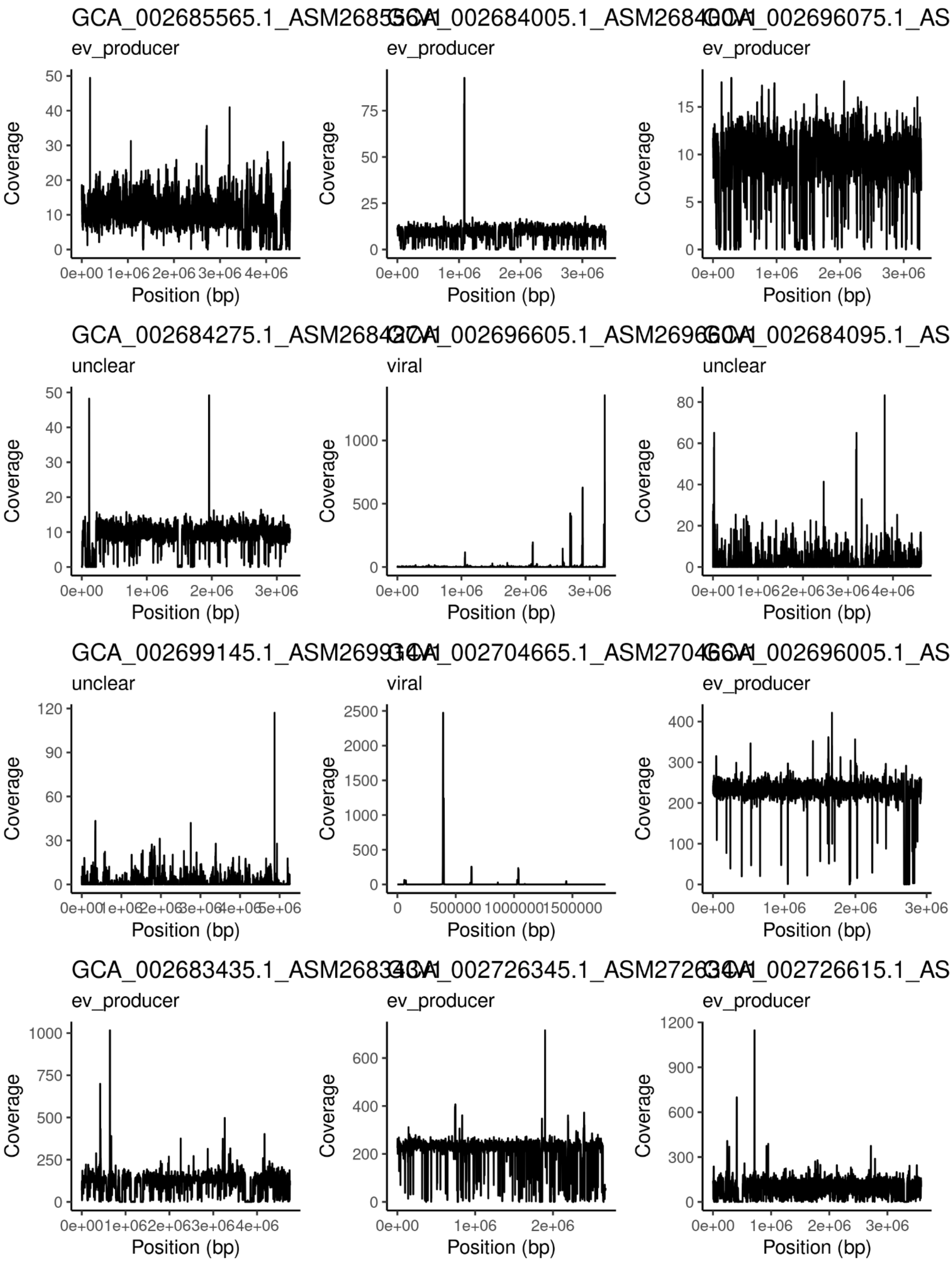
GCA_002685565.1_ASM268556
ev_producer
GCA_002684005.1_ASM268400
ev_producer
GCA_002696075.1_AS
ev_producer
GCA_002684275.1_ASM268427
unclear
GCA_002696605.1_ASM269660
viral
GCA_002684095.1_AS
unclear
GCA_002699145.1_ASM269914
unclear
GCA_002704665.1_ASM270466
viral
GCA_002696005.1_AS
ev_producer
GCA_002683435.1_ASM268343
ev_producer
GCA_002726345.1_ASM272634
ev_producer
GCA_002726615.1_AS
ev_producer
Coverage
Position (bp)

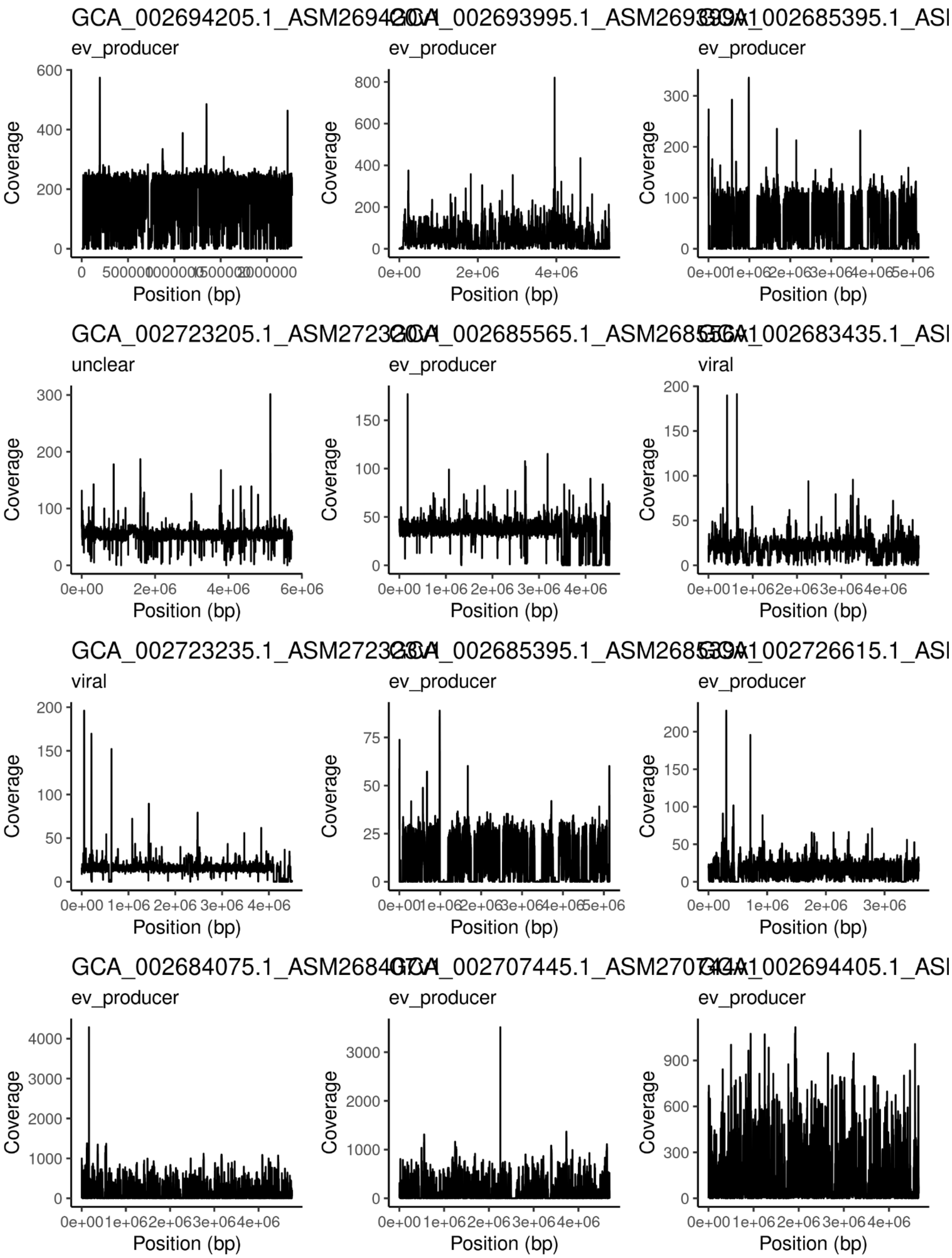

GCA_002694205.1_ASM269420
ev_producer
GCA_002693995.1_ASM269399
ev_producer
GCA_002685395.1_ASI
ev_producer
GCA_002723205.1_ASM272320
unclear
GCA_002685565.1_ASM268556
ev_producer
GCA_002683435.1_ASI
viral
GCA_002723235.1_ASM272323
viral
GCA_002685395.1_ASM268539
ev_producer
GCA_002726615.1_ASI
ev_producer
GCA_002684075.1_ASM268407
ev_producer
GCA_002707445.1_ASM270744
ev_producer
GCA_002694405.1_ASI
ev_producer
Coverage
Position (bp)

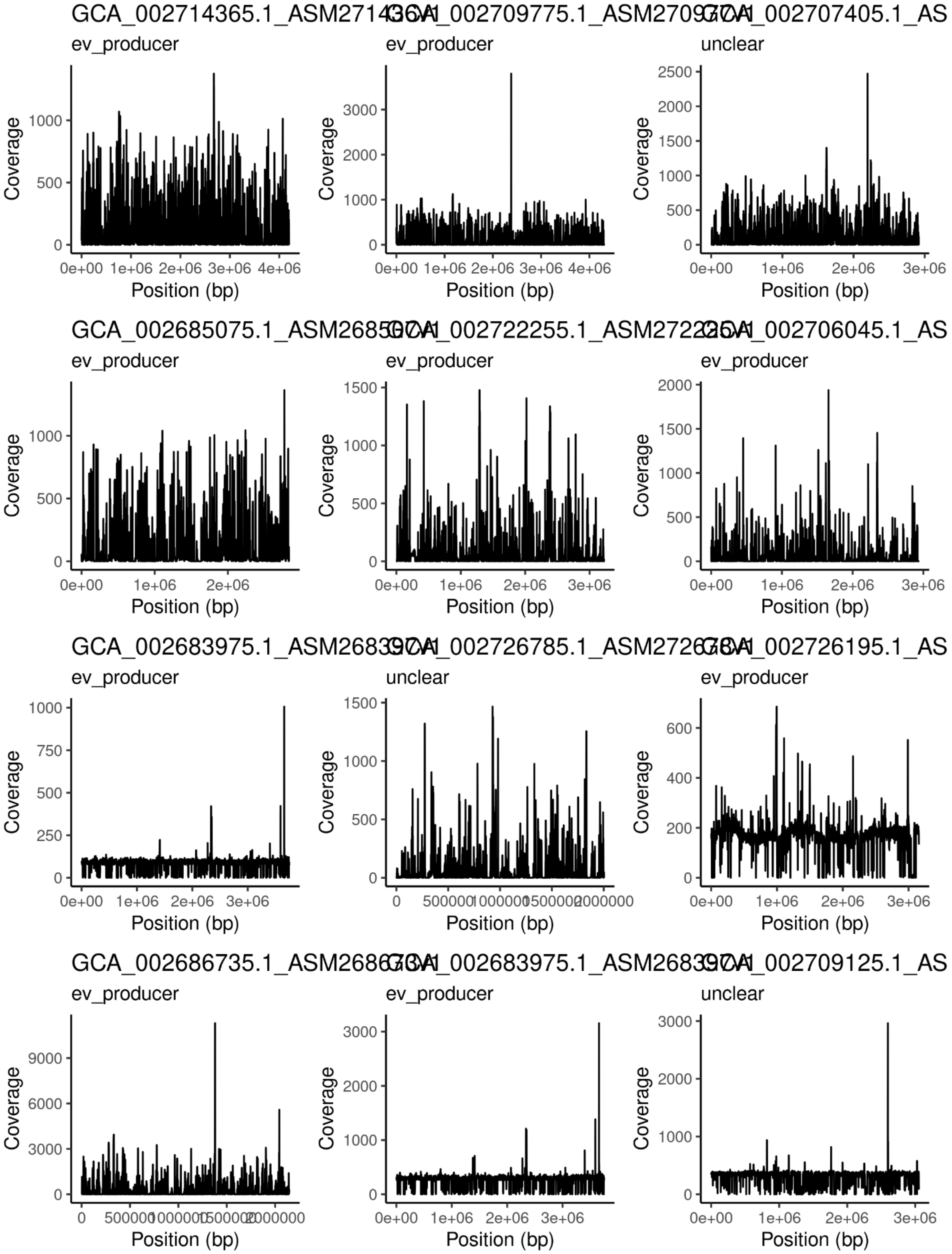
GCA_002714365.1_ASM271436v1
ev_producer
GCA_002709775.1_ASM270977v1
ev_producer
GCA_002707405.1_AS
unclear
GCA_002685075.1_ASM268507v1
ev_producer
GCA_002722255.1_ASM272225v1
ev_producer
GCA_002706045.1_AS
ev_producer
GCA_002683975.1_ASM268397v1
ev_producer
GCA_002726785.1_ASM272678v1
unclear
GCA_002726195.1_AS
ev_producer
GCA_002686735.1_ASM268673v1
ev_producer
GCA_002683975.1_ASM268397v1
ev_producer
GCA_002709125.1_AS
unclear
Coverage
Position (bp)

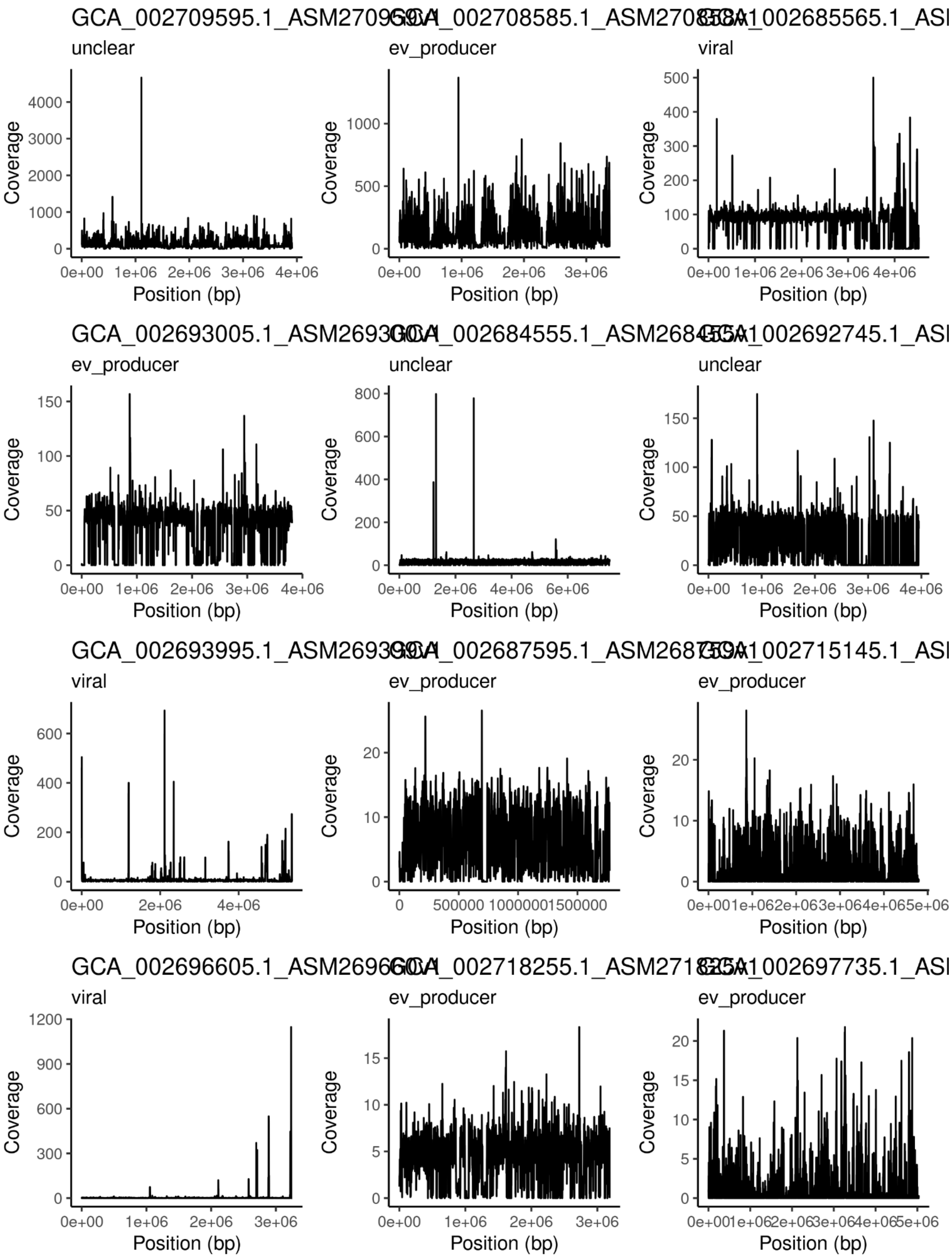
GCA_002709595.1_ASM270959
unclear
GCA_002708585.1_ASM270858
ev_producer
GCA_002685565.1_ASI
viral
GCA_002693005.1_ASM269300
ev_producer
GCA_002684555.1_ASM268455
unclear
GCA_002692745.1_ASI
unclear
GCA_002693995.1_ASM269399
viral
GCA_002687595.1_ASM268759
ev_producer
GCA_002715145.1_ASI
ev_producer
GCA_002696605.1_ASM269660
viral
GCA_002718255.1_ASM271825
ev_producer
GCA_002697735.1_ASI
ev_producer
Coverage
Position (bp)

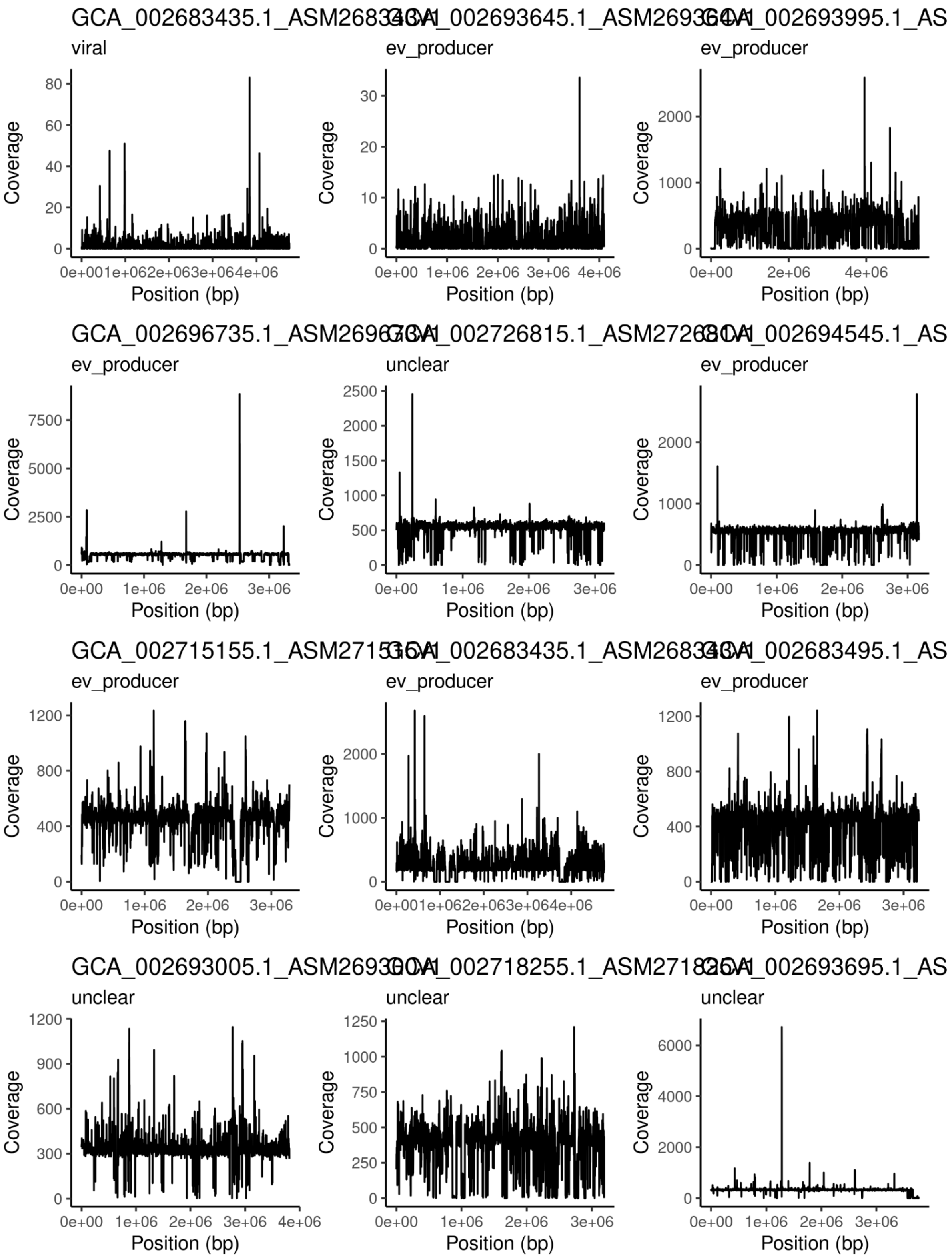
GCA_002683435.1_ASM268343v1
viral
GCA_002693645.1_ASM269364v1
ev_producer
GCA_002693995.1_AS
ev_producer
GCA_002696735.1_ASM269673v1
ev_producer
GCA_002726815.1_ASM272681v1
unclear
GCA_002694545.1_AS
ev_producer
GCA_002715155.1_ASM271515v1
ev_producer
GCA_002683435.1_ASM268343v1
ev_producer
GCA_002683495.1_AS
ev_producer
GCA_002693005.1_ASM269300v1
unclear
GCA_002718255.1_ASM271825v1
unclear
GCA_002693695.1_AS
unclear
Coverage
Position (bp)

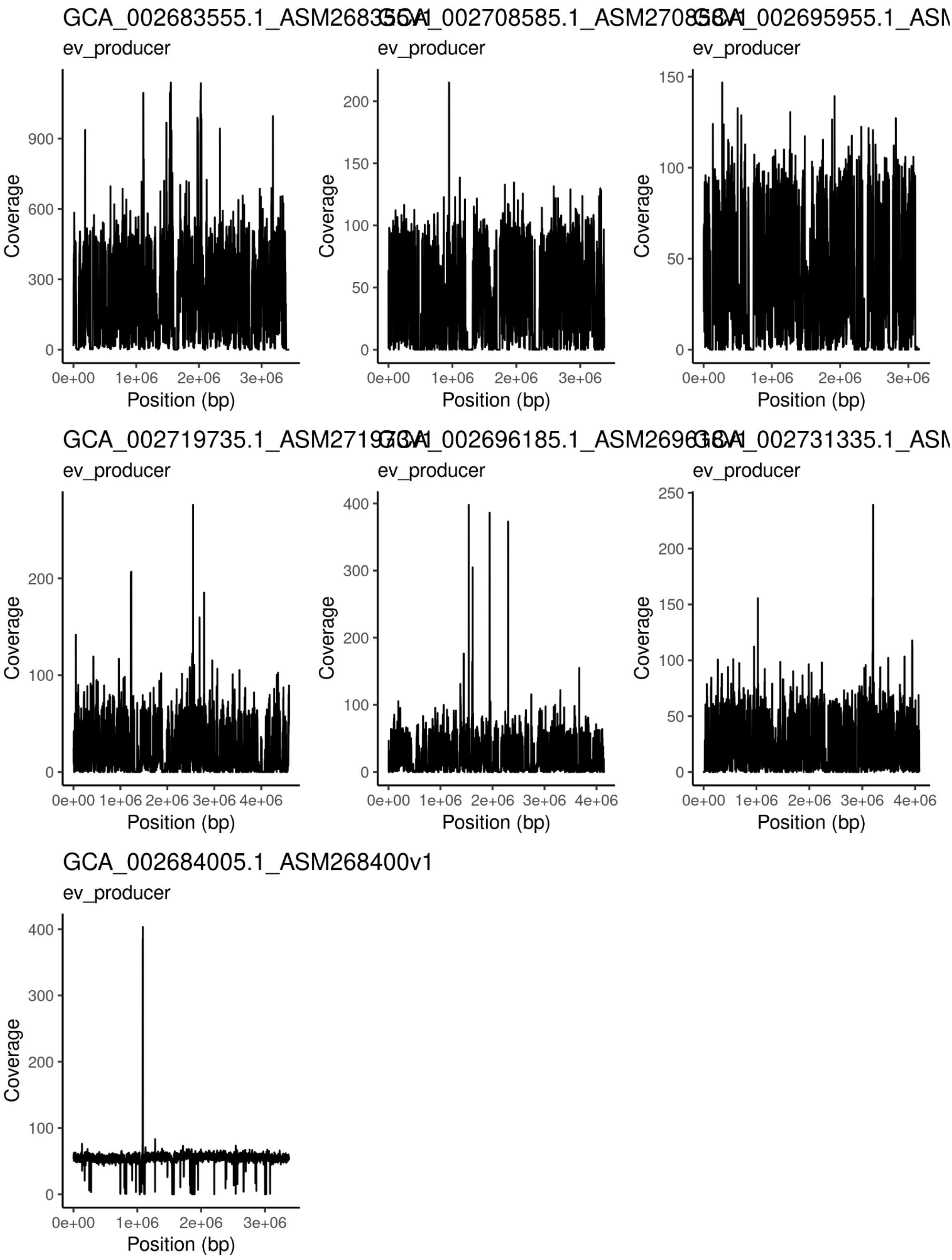

ev_producer
Coverage
Position (bp)
GCA_002684005.1_ASM268400v1
